# Multimodal and spatially resolved profiling identifies distinct patterns of T-cell infiltration in nodal B-cell lymphoma entities

**DOI:** 10.1101/2022.11.04.514366

**Authors:** Tobias Roider, Marc A. Baertsch, Donnacha Fitzgerald, Harald Voehringer, Berit J. Brinkmann, Felix Czernilofsky, Mareike Knoll, Laura Llaó-Cid, Peter-Martin Bruch, Nora Liebers, Christian M. Schürch, Verena Passerini, Alexander Brobeil, Gunhild Mechtersheimer, Carsten Müller-Tidow, Oliver Weigert, Martina Seiffert, Garry P. Nolan, Wolfgang Huber, Sascha Dietrich

**Affiliations:** European Molecular Biology Laboratory (EMBL), Heidelberg, Germany; Molecular Medicine Partnership Unit (MMPU), Heidelberg, Germany; Department of Medicine V, Hematology, Oncology and Rheumatology, University of Heidelberg, Heidelberg, Germany; Department of Microbiology and Immunology, Stanford University School of Medicine, Stanford, CA, USA; Clinical Cooperation Unit Molecular Hematology/Oncology, German Cancer Research Center (DKFZ), Heidelberg, Germany; Division of Molecular Genetics, German Cancer Research Center (DKFZ), Heidelberg, Germany; Department of Hematology and Oncology, University Hospital Düsseldorf, Düsseldorf, Germany; National Center for Tumor Diseases (NCT), Heidelberg, Germany; German Cancer Research Center (DKFZ), Heidelberg, Germany; Department of Pathology and Neuropathology, University Hospital and Comprehensive Cancer Center Tübingen, Tübingen, Germany; Department of Medicine III, Laboratory for Experimental Leukemia and Lymphoma Research (ELLF), Ludwig-Maximilians-University (LMU) Hospital, Munich, Germany; Department of Pathology, University of Heidelberg, Heidelberg, Germany; German Cancer Consortium (DKTK), Munich, Germany; Department of Pathology, Stanford University School of Medicine, Stanford, CA, USA

**Author notes:** Correspondence (T.R.), (S.D.). These first authors contributed equally (Tobias Roider, Marc A. Baertsch). These senior authors contributed equally (Wolfgang Huber, Sascha Dietrich).

## Abstract

T-cell-engaging immunotherapies have improved the treatment of nodal B-cell lymphoma, but responses vary highly. Future improvements of such therapies require better understanding of the variety of lymphoma-infiltrating T-cells. We employed single-cell RNA and T-cell receptor sequencing alongside quantification of surface proteins, flow cytometry and multiplexed immunofluorescence on 101 lymph nodes from healthy controls, and patients with diffuse large B-cell, mantle cell, follicular, or marginal zone lymphoma. This multimodal resource revealed entity-specific quantitative and spatial aberrations of the T-cell microenvironment. Clonal PD1^+^ *TCF7*^-^ but not PD1^+^ *TCF7*^+^ cytotoxic T-cells converged into terminally exhausted T-cells, the proportions of which were variable across entities and linked to inferior prognosis. In follicular and marginal zone lymphoma, we observed expansion of follicular helper and IKZF3^+^ regulatory T-cells, which were clonally related and inversely associated with tumor grading. Overall, we portray lymphoma-infiltrating T-cells with unprecedented comprehensiveness and decipher both beneficial and adverse dimensions of T-cell response.

## Introduction

Nodal B-cell non-Hodgkin lymphomas (B-NHL) represent a heterogeneous group of indolent and aggressive malignancies that grow mainly in the lymph node (LN). Extensive genetic and transcriptomic profiling has revealed disease-specific mutational signatures and pathway dependencies, paving the way for targeted molecular therapy [1-8]. However, in recent years, T-cell-engaging immunotherapies, such as bispecific antibodies or chimeric antigen receptor (CAR) T-cells, have emerged among the leading treatment options for refractory and relapsed B-NHL patients [9-12]. Tailoring these treatment approaches to different B-NHL entities and identifying the vulnerabilities of their microenvironment requires systematic investigation of the variety and functions of tumor-infiltrating T-cells – analogously to studying the genetic and transcriptomic makeup of tumor cells as a prerequisite for tailoring targeted therapies.

Traditionally, T-cell phenotyping studies have been based on immunohistochemistry or flow cytometry, and have established a fundamental understanding of the T-cell microenvironment in nodal B-NHL [13]. In recent years, single-cell RNA sequencing (scRNA-seq) emerged as a powerful tool to capture the heterogeneity of tumor-infiltrating T-cells and consequently became an integral part of T-cell phenotyping efforts [14, 15]. We and others have pioneered the investigation of transcriptional heterogeneity of LN-derived T-cells in B-NHL, but with the limitation of low sample sizes or having focused only on follicular lymphoma (FL, indolent) [16-18]. Stand-alone scRNA-seq studies additionally face the problem to align gene expression profiles with known T-cell subsets that have been defined for decades based on surface epitopes and transcription factors [19].

Here, we employed cellular indexing of transcriptomes and epitopes by sequencing (CITE-seq), which simultaneously captures transcript and surface epitope abundances at single-cell resolution, and thus enables a multimodal identification of T-cell phenotypes [20, 21]. Moreover, we took a substantially wider view by collecting more than 100 LN patient samples and by including, besides FL, other B-NHL entities with only little or no prior groundwork: diffuse large B-cell lymphoma (DLBCL, aggressive), marginal zone lymphoma (MZL, indolent), and mantle cell lymphoma (MCL, mixed). We identified and quantitated fine-grained T-cell subsets and determined their clonality using full-length single-cell T-cell receptor sequencing. We further assessed the ability of multicolor flow cytometry and highly multiplexed immunofluorescence to reproduce the quantification of these T-cell subsets and to localize them within the tumor microenvironment. Based on these data, we created a spatially resolved reference map for LN-derived T-cells in nodal B-NHL and identified entity-specific patterns of T-cell infiltration.

## Results

### Study and sample overview

We collected 101 LN samples (Supplementary Figure 1A) from patients with DLBCL (n = 28), FL (n = 30), MZL (n = 15), MCL (n = 15), or from patients without evidence of malignancy (tumor-free / reactive LN [rLN], n = 13). LN samples from 38 patients were collected at initial diagnosis, while 50 LN samples were collected from patients who had received one or more prior lines of systemic treatment (Supplementary Figure 1A). To minimize potential effects on T-cell infiltration patterns, relapse samples were collected at least 3 months after cessation of the preceding systemic treatment. T-cell proportions of malignant LN samples determined by flow cytometry showed a broad variation (Supplementary Figure 1A) but were not significantly associated with pre-treatment status, sex, age, or B-NHL entity (Supplementary Figure 1B-E). Patient characteristics are summarized in Supplementary Table 1.

### Lymph node-derived T-cells can be divided into fourteen multimodally defined subsets

We used CITE-seq, a droplet-based single-cell sequencing technology that measures the transcriptome and selected cell surface proteins, to profile T-cells from 51 LN patient samples. Surface proteins were detected using 70 oligonucleotide-tagged antibodies (Supplementary Table 2). After quality control and in-silico sorting, we obtained data for 74,112 T-cells with a median of 1,190 T-cells per patient sample. Unsupervised clustering based on a weighted combination of transcriptome and epitope similarities (weighted-nearest-neighbor, WNN) [22] grouped the T-cells into proliferating (T_Pr_), conventional helper (T_H_) and follicular helper (T_FH_), regulatory (T_REG_), cytotoxic (T_TOX_), and double negative T-cells (T_DN_, Figure 1A). Differentially expressed genes and proteins (Figure 1B, C) were associated with lineage (CD4, CD8), functional specialization (e.g., *FOXP3, ASCL2*), cytotoxicity (e.g., *GZMA, GZMK*) or proliferation (e.g., *MKI67*). These groups could be further partitioned into CD4^+^ and CD8^+^ naïve T-cells, central memory (CM_1_, CM_2_) T_H_ cells, central memory (CM_1_, CM_2_) and effector memory (EM_1_, EM_2_) T_REG_ cells, and effector memory (EM_1_, EM_2_, EM_3_) T_TOX_ cells. At this level of granularity, differentially expressed markers (Figure 1B, C) were linked to differentiation (e.g., CD45RA, CD45RO, CD62L), homing and migration (e.g., *KLF2, CCR7*), activation (e.g., CD69, CD38, CD278), and inhibition (e.g., PD1, TIM3, LAG3). This high-granularity classification was supported by a gene regulatory network analysis [23], which highlighted differential activities of specific transcription factors (e.g., KLF2 [24], TCF7 [25], FoxP3 [26], ASCL2 [27], Figure 1D). Based on that, we compiled profiles of the most discriminating and biologically interpretable surface proteins, genes and transcription factors that facilitate the distinction of all fourteen T-cell subsets (Figure 1E). Further extended profiles, for instance to identify T-cell subsets in scRNA-seq data using previously published scoring approaches [28], are provided in Supplementary Table 3.

**Figure 1.**
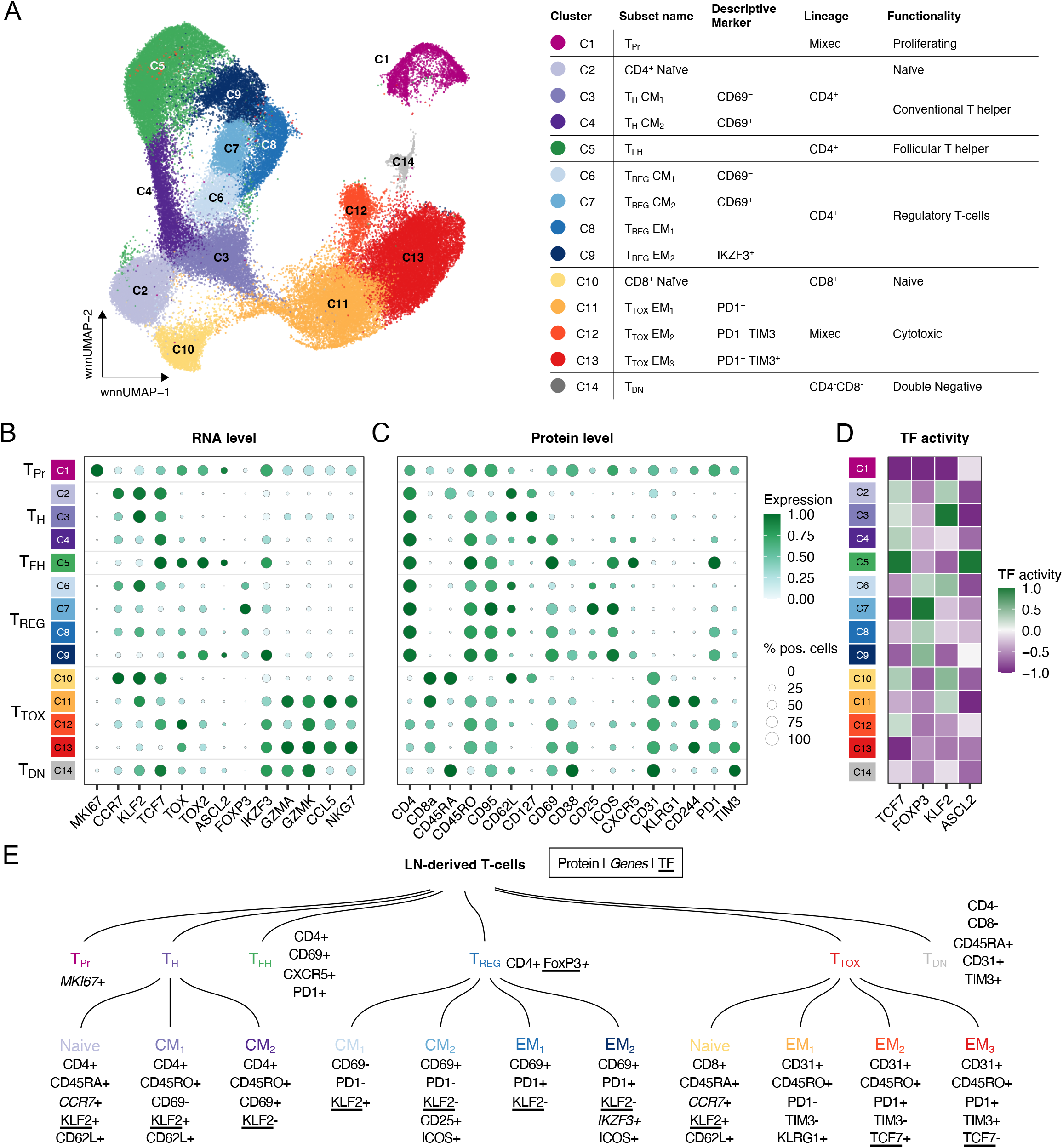
Lymph node-derived T-cells can be divided into fourteen multimodally defined subsets. A) CITE-seq data from 51 primary LN patient samples were integrated and jointly visualized using UMAP. Cells were colored with respect to their cluster based on a shared nearest neighbor-based algorithm. The adjacent table summarizes all clusters including used subset names, lineages and functionality. B and C) Dot plot showing the expression of important marker genes and proteins to identify all T-cell subsets. Size and color of the dots indicate the percentage of positive cells and scaled gene / protein expression, respectively. Values were scaled between 0 and 1. Heatmap showing the inferred activity of selected TF, as indicated. Y axis is identical to panel B and C. Values were scaled between -1 and 1. Dendrogram summarizing the 14 multimodally defined T-cell subsets including most important and interpretable marker genes, proteins, and TF. CM: Central memory. EM: Effector memory. LN: Lymph node. TF: Transcription factor.

### Multicolor flow cytometry reproduces multimodally defined T-cell subsets

Having used CITE-seq to discover and characterize the T-cell subsets, we employed flow cytometry to enable their rapid and cost-effective quantification in large sample numbers. We built gradient boosting classifiers [29] to identify the most discriminatory surface markers between multimodally defined subsets (Figure 2A). While this yielded accurate results for most T-cell subsets (Supplementary Figure 2A), the distinction among T_REG_ and the detection of T_Pr_ cells could be improved by additional intracellular markers that were selected based on their accessibility by flow cytometry and the signature profiles above (Ki67, FoxP3, IKZF3, Supplementary Figure 2B). After removal of redundant (e.g., CD95, CD127) and less important markers, we thus compiled a 12- and 13-plex flow cytometry panel (Supplementary Table 4) and established gating strategies supported by the hypergate algorithm [30], which enabled classification of all 14 multimodally defined T-cells subsets (Supplementary Figures 2C, 3). We correlated the subset proportions determined by CITE-seq and flow cytometry across a total of 13 LN samples and observed a median Pearson coefficient of 0.92 across all subsets (Figure 2B). We then applied these panels to an independent cohort of 50 LN samples, which was then used for further quantitative analysis of T-cell infiltration patterns.

**Figure 2.**
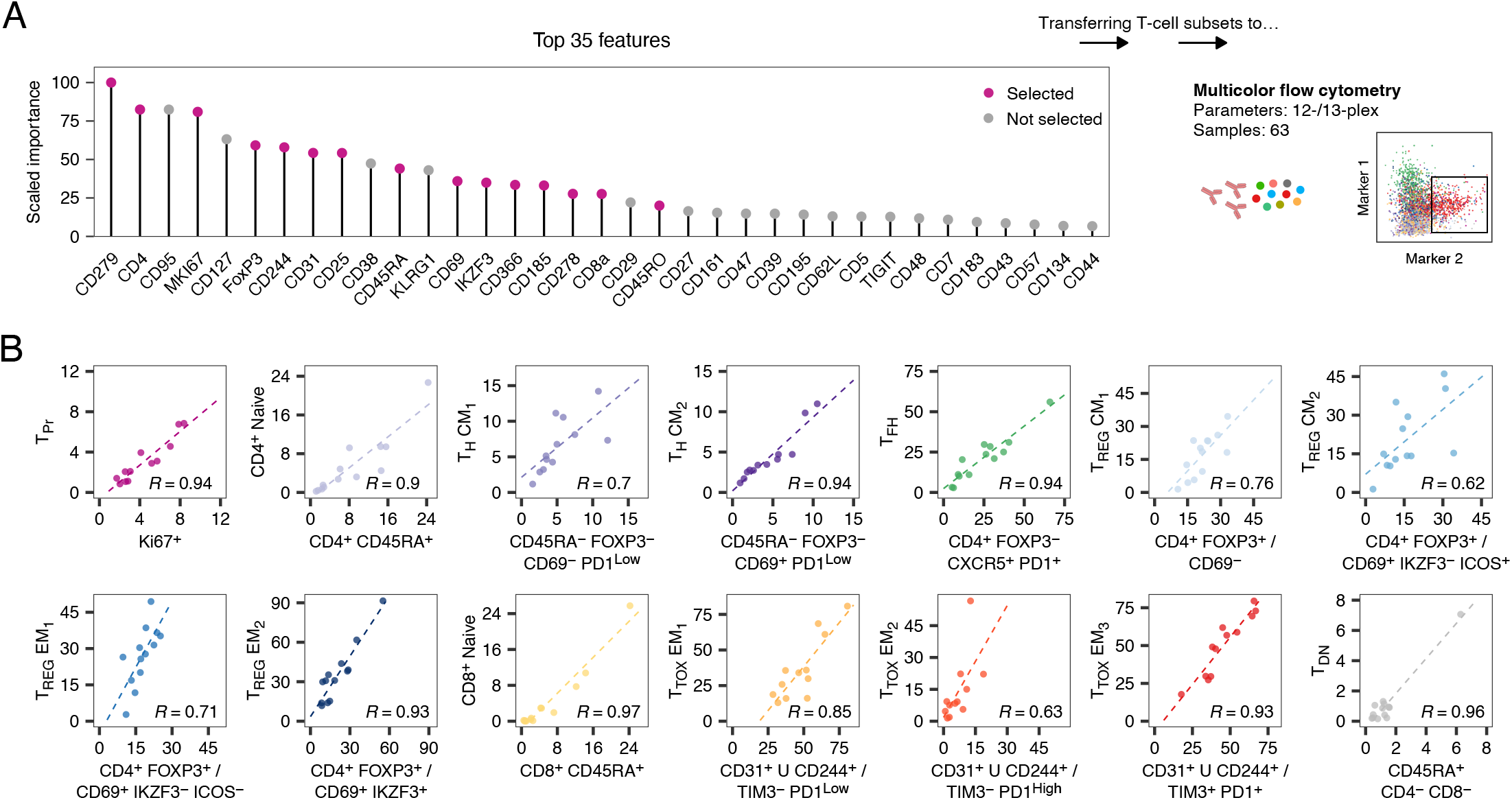
Multicolor flow cytometry reproduces multimodally defined T-cell subsets. A) Most important features to distinguish multimodally defined T-cell subsets using a gradient boosting classifier. Only features that are routinely accessible by flow cytometry were considered for the model. B) Percentages of all T-cell subsets determined by flow cytometry (x axis) and CITE-seq (y axis) were correlated for n = 13 biologically independent samples. X axis title indicates the applied gating strategy. The symbol U indicates merging of two populations. Pearson’s correlation coefficient is given for each panel (R).

### Nodal B-NHL entities have characteristic quantitative patterns of T-cell infiltration

To provide a systematic and high-resolution analysis of the T-cell microenvironment in B-NHL, we combined CITE-seq and flow cytometry data, determined the proportion of each subset per sample, and compared each B-NHL entity with tumor-free LN samples (Figure 3A). B-NHL LN were characterized by a lack of CD69^-^ CM_1_ and CD69^+^ CM_2_ T_H_ cells, and CD4^+^ / CD8^+^ naive T-cells (Figure 3A). Conversely, PD1^+^ TIM3^-^ T_TOX_ EM_2_ and PD1^+^ TIM3^+^ EM_3_ T_TOX_ cells, T_Pr_ cells, and CD69^+^ T_REG_ CM_2_ cells were significantly enriched in B-NHL LN (Figure 3A). FL and MZL were additionally characterized by significant enrichment of T_FH_ and IKZF3^+^ T_REG_ EM_2_ cells (Figure 3A). We also noted a significant increase of CD69^+^ T_REG_ CM_2_ cells in MCL, FL, and MZL, whereas T_FH_ cells were depleted in DLBCL (Figure 3A). To gain a broader overview of these differences across all B-NHL entities, we used principal component analysis (PCA) on the table of subset proportions (Figure 3B). Based on the first two principal components (PC), we identified three major groups (I-III) represented by tumor-free (I), DLBCL and MCL (II), and FL and MZL LN (III, Figure 3B). PC1 (Figure 3C) and PC2 (Figure 3D) had high loadings on the characteristic T-cell subsets highlighted in Figure 3A: CD4^+^ and CD8^+^ naïve T-cells, and PD1^+^ TIM3^+^ T_TOX_ EM_3_ cells (PC1), and T_FH_ and IKZF3^+^ T_REG_ EM_2_ cells (PC2).

**Figure 3.**
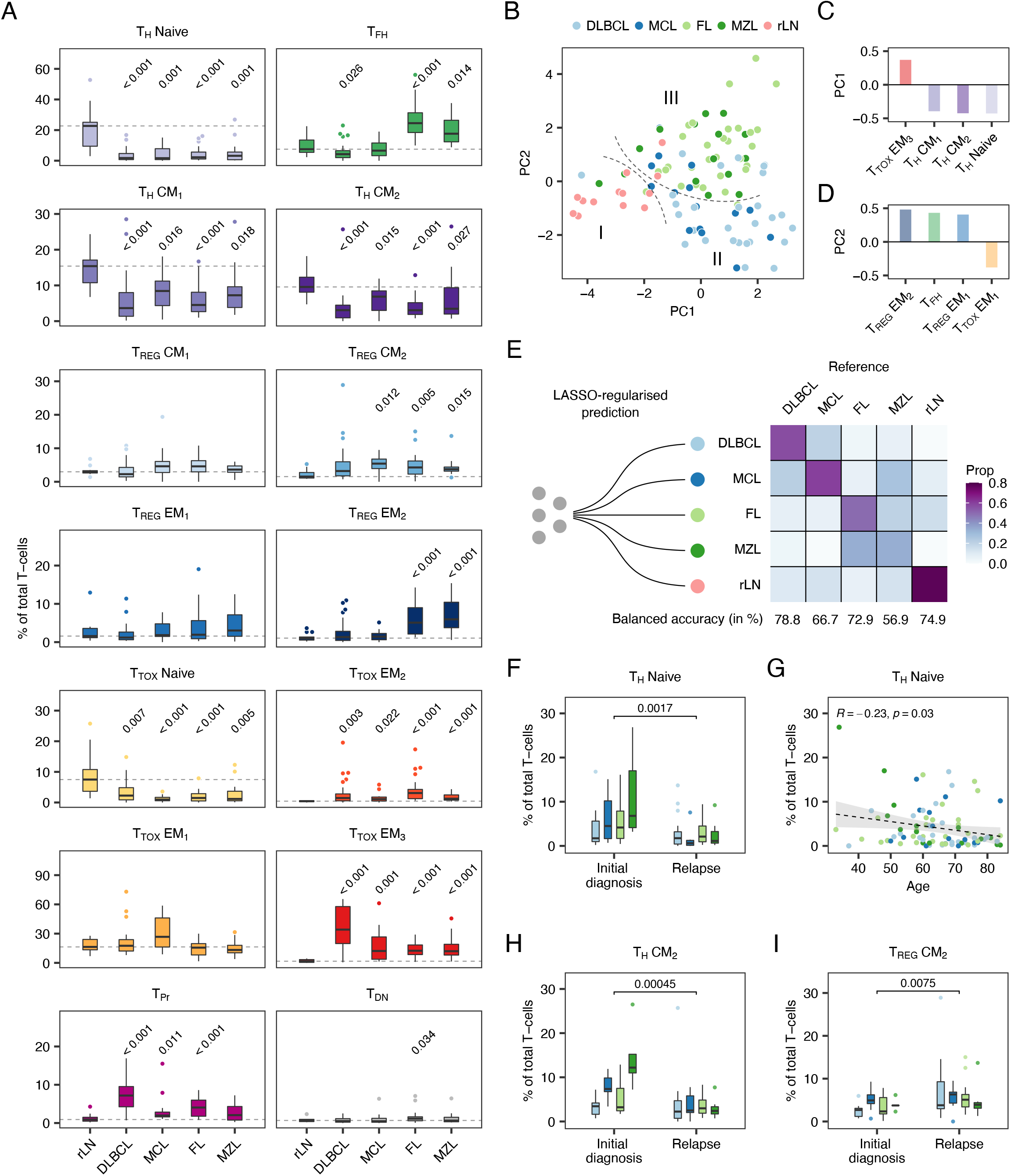
Nodal B-NHL entities have characteristic quantitative patterns of T-cell infiltration. A) T-cell subset proportions determined by CITE-seq or flow cytometry are illustrated in box plots (n = 101). Outliers are shown as individual dots. Each entity and subset were tested versus tumor-free samples (rLN) using the Wilcoxon-test. P values were corrected for multiple testing using the Benjamini-Hochberg procedure. Dashed lines indicate the median of rLN. B-D) Principal component analysis based on the subset and overall T-cell proportions (B) including the top four loadings of principal component 1 (C) and 2 (D) are shown. Dashed lines (B) highlight three groups (I-III) of samples. E) Confusion matrix based on a LASSO-regularized multinomial logistic regression model and estimated classification accuracy using leaving-one-out cross validation based on subset and overall T-cell proportions. F-I) Patient characteristics were evaluated in a multivariate model regarding their impact on the proportions of all 14 T-cell subsets. Shown are the four most significant associations. P values and/or correlation coefficients were calculated using unpaired Wilcoxon-test (F, H, I) or Pearson’s linear correlation (G). Box plots or dots are colored by entity as in panel B and E. PC: Principal component.

To further explore to what extent T-cell composition is distinctive for different B-NHL entities and tumor-free LN, we built classifiers using LASSO-regularized multinomial logistic regression and estimated classification accuracy using nested leave-one-out cross-validation (Figure 3E, Supplementary Figure 4A). Accuracy was best for distinguishing between tumor-free and malignant LN (balanced accuracy of 74.5 %); moreover, DLBCL and MCL could be differentiated with similar accuracy (Figure 3E). A third, well distinguishable group was formed by FL and MZL (Figure 3E). These results indicate that different entities have distinct patterns of T-cell infiltration (Supplementary Figure 4A), though – based on our current data – classification does not provide diagnostic accuracy, which might be the consequence of inter- and/or intra-tumor heterogeneity [16].

To explore the potential role of patient-inherent characteristics, we fit multivariate linear models using sex, age, treatment status, and cell-of-origin (only DLBCL) as covariates, and the proportion of each T-cell subset as dependent variable (Supplementary Figure 4B). We found that pre-treatment (Figure 3G, p < 0.001) and higher age (Figure 3H, p = 0.04) were associated with a lower proportion of naïve CD4^+^ T-cells. Pre-treatment was also linked to a lower proportion of CD69^+^ T_H_ CM_2_ cells (Figure 3I, p < 0.001) and a higher proportion of CD69^+^ T_REG_ CM_2_ cells (Figure 3J, p < 0.001), while we observed no statistically significant impact on the T-cell composition for sex and cell-of-origin (Supplementary Figure 4B). Larger sample sizes might be necessary to detect less strong associations; but overall, patient characteristics had only a moderate impact on the T-cell composition compared to entity-specific differences.

### Entity-specific T-cell compositions result from differential clonal expansion of CD4^+^ and CD8^+^ T-cell subsets

To investigate if the enrichment of T-cell subsets results from their clonal expansion, we performed full-length single T-cell receptor (scTCR) sequencing alongside 5’ scRNA-seq in a representative subset of patient samples (n = 11, Supplementary Figure 1A, Supplementary Table 1). After quality control, 5’ scRNA and full-length TCR data were available for 30,198 T-cells with a median of 2,892 cells per patient sample. We mapped the 5’ scRNA onto the CITE-seq reference data above, indicating high consistency of the inferred subset proportions between both modalities with a median correlation coefficient of R = 0.92 (Supplementary Figure 5A). Then, we compiled the scTCR data to clonotypes based on their complementarity-determining regions (CDRs) [31] and projected them onto the reference UMAP (Figure 4A). We found that clonally expanded T_TOX_ EM_1_ cells were present across all entities and in tumor-free LN (Figure 4A), while clonally expanded PD1^+^ TIM3^+^ T_TOX_ EM_3_ were limited to DLBCL, FL and MZL (Figure 4A). Clonality of T_TOX_ cells was not restricted to *CD8*^+^ but included – albeit to lower extent – also *CD4*^*+*^ T_TOX_ cells (Supplementary Figure 5B-C). In addition, T_FH_ and IKZF3^+^ T_REG_ EM_2_ cells were clonally expanded exclusively in FL and MZL (Figure 4A). Consequently, the TCR diversity was substantially reduced in DLBCL, FL, and MZL samples compared to tumor-free and MCL samples (Supplementary Figure 5D-F).

**Figure 4.**
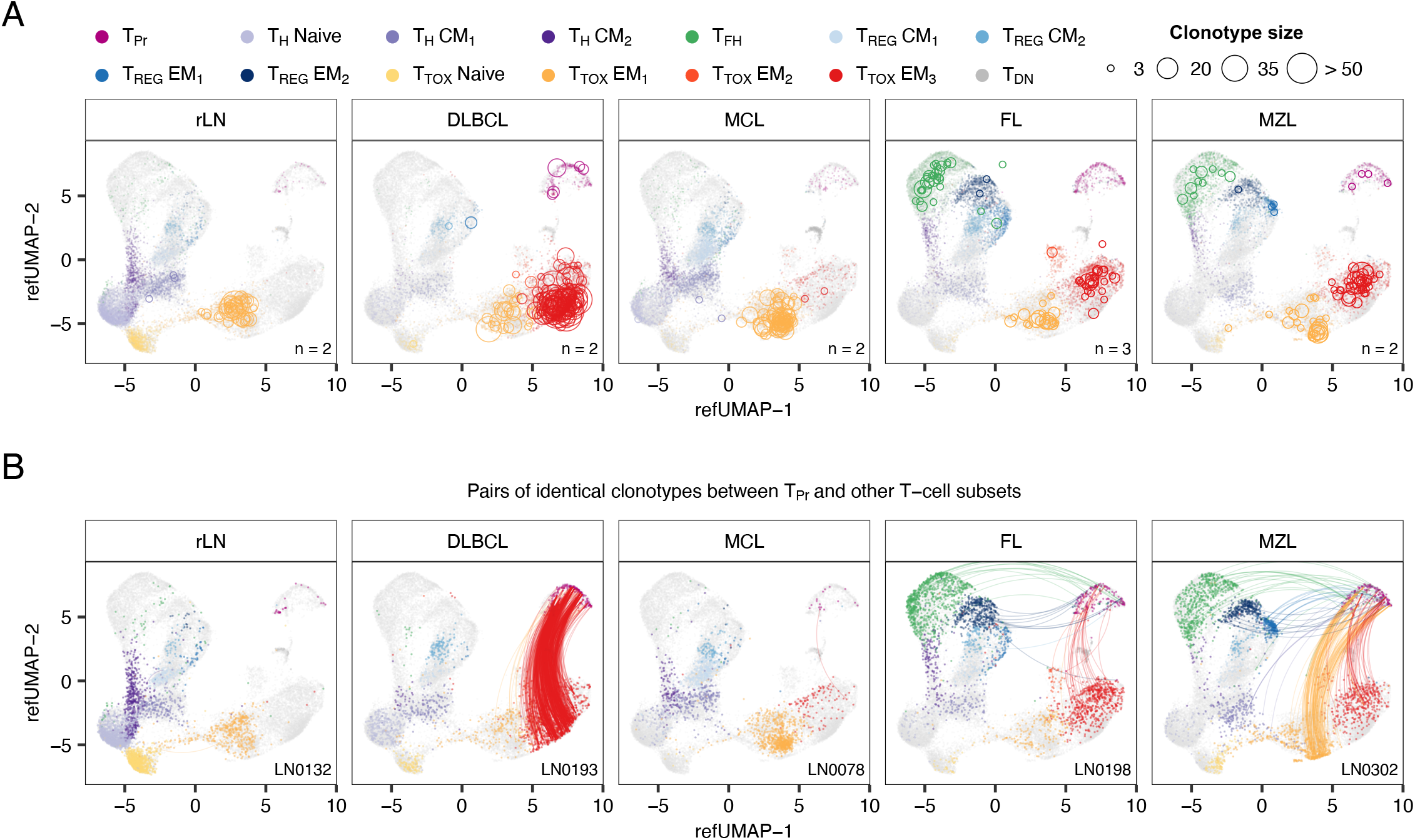
Entity-specific T-cell compositions result from differential clonal expansion of CD4+ and CD8+ T-cell subsets. 5’ scRNA alongside full-length TCR repertoire data were mapped to the CITE-seq reference dataset. In grey, all cells with 5’ scRNA data are shown, whereas colored cells belong to samples derived from specific entities (A) or samples (B), as indicated. A) Circles represent the number of cells with identical TCR clonotype within the same subpopulation. B) Shown are mapped cells from five representative samples. Lines connect all proliferating cells with any other cell given that both have identical TCR clonotypes. TCR: T-cell receptor.

We further used scTCR data to track the original identity of T_Pr_ cells, that is at the time of sample collection temporarily masked by an S or G_2_M phase-dependent gene expression signature [32]. Apart from MCL and tumor-free LN, which both harbored very low proportions of T_Pr_ cells (Figure 4A, B), we identified groups of shared clonotypes in DLBCL, FL, and MZL predominantly between T_Pr_ and PD1^+^ TIM3^+^ T_TOX_ EM_3_ cells (Figure 4B). To a lower extent, TCR clonotypes were also shared between T_Pr_ and T_FH_ cells, and between T_Pr_ and IKZF3^+^ T_REG_ EM_2_ cells in FL and MZL (Figure 4B). Overall, this analysis highlights that altered T-cell microenvironment is a result from active proliferation and differential expansion of specific T-cell subsets.

### PD1^+^ TCF7^-^ T_TOX_ cells converge into terminally exhausted T-cells with variable proportions within and across entities

Ongoing antigen exposure and clonal expansion, as in cancer or chronic infection, can ultimately result in T-cell exhaustion and loss of anti-tumor activity [33]. To unveil the process of T-cell exhaustion across B-NHL entities in more detail, we performed a trajectory analysis [34] of T_TOX_ cells based on the CITE-seq data. We identified two paths: (I) one from naïve to PD1^+^ TIM3^-^ T_TOX_ EM_2_ cells, and (II) another one from naïve to PD1^+^ TIM3^+^ T_TOX_ EM_3_ cells (Figure 5A). Apart from TIM3, both expression (Figure 5B) and inferred transcription factor activity [23] of *TCF7* was an important discriminator between trajectory I and II (Figure 1D). TCF1 (encoded by *TCF7*) is a hallmark of stemness and longevity and its presence (trajectory I) or absence (trajectory II) indicates maintained or impaired self-renewal capacity, respectively [35], thereby suggesting that only trajectory II converges into terminally exhausted T-cells.

**Figure 5.**
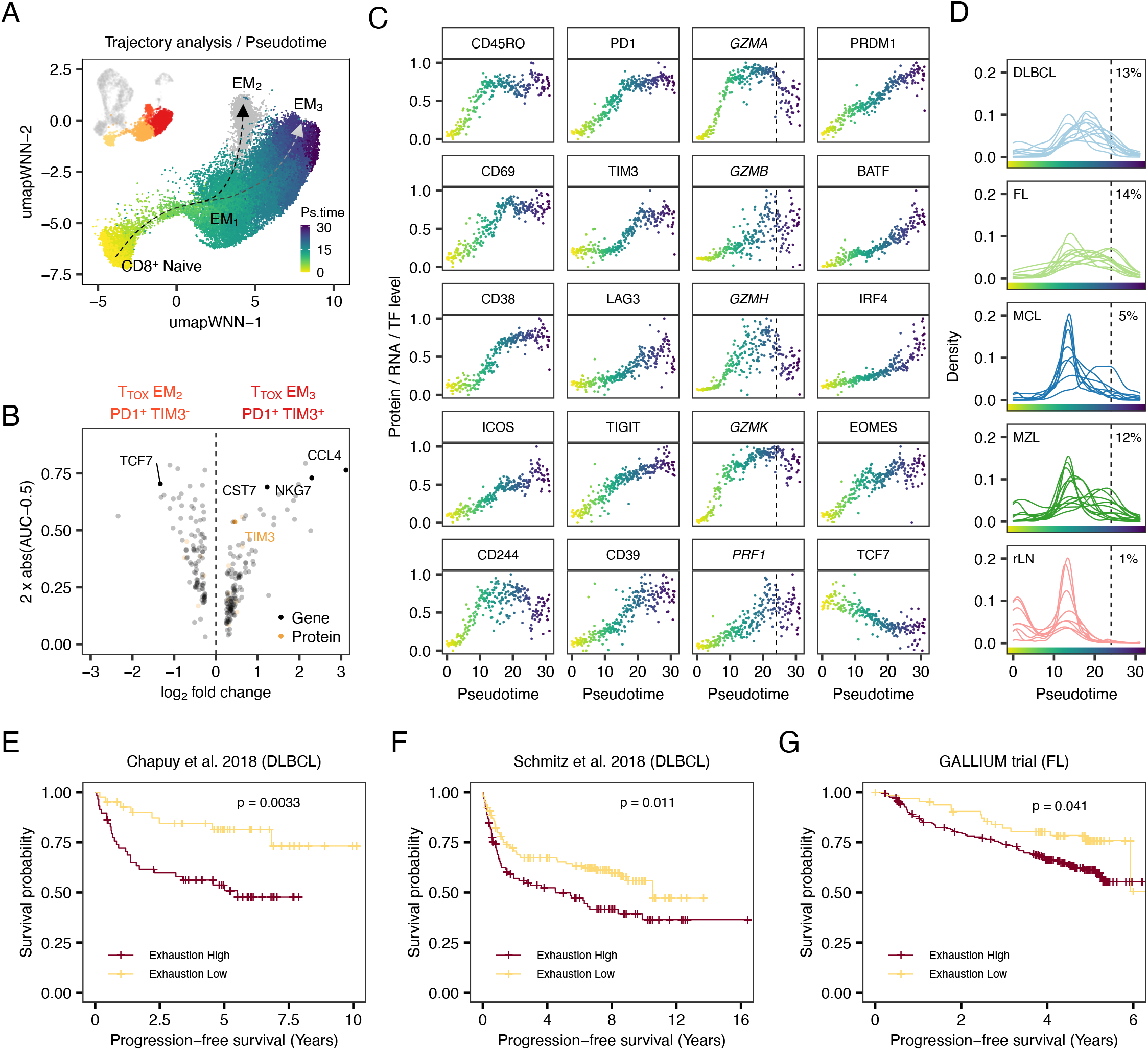
PD1+ TCF7− cytotoxic T-cells converge to terminally exhausted T-cells with variable proportions within and across entities. A) Combined trajectory and pseudotime analysis were performed using CITE-seq expression profiles of TTOX cells starting from naive CD8^+^ T-cells. Arrows illustrate trajectories, while cells are colored by pseudotime. B) Volcano plot illustrating differentially expressed genes and proteins between PD1^+^ TIM3^+^ TTOX EM3 and PD1^+^ TIM3^-^ TTOX EM2 cells. C) Protein expression (first and second column), gene expression (third column) or inferred TF activity (fourth column) are illustrated along binned pseudotime, as shown in panel A. Values were scaled between 0 and 1. Dashed line in *GZMA* plot indicates threshold when T-cells were considered terminally exhausted. D) Shown is the density of cells for each single patient along pseudotime. Number indicates median percentage of terminally exhausted T-cells across all LN patient samples for each entity. E-G) Bulk RNA-seq data from DLBCL (E, F) and FL (G) patients were deconvoluted based on a gene expression signature of terminally exhausted T-cells. Kaplan-Meier plots with p values of corresponding log-rank test. LN: Lymph node. TF: Transcription factor.

To further resolve gradual changes along trajectory II, we applied pseudotime analysis [34] and ranked the cells starting from naive T_TOX_ cells (Figure 5A). We found that pseudotime was strongly linked to a continuously increasing expression of both differentiation and activation markers, such as CD45RO, CD69, CD38, ICOS, and inhibitory molecules, such as PD1, TIM3, LAG3, TIGIT and CD39 (Figure 5C). Cells with the highest levels of inhibitory receptors had reduced RNA expression levels of effector molecules, particularly of *GZMA* and *GZMH*, and reduced protein levels of CD244, which mediates non-MHC cytotoxicity [36] (Figure 5C). Likewise, the inferred activity [23] of the transcription factors PRDM1, BATF, IRF4 and EOMES, which have previously been associated with T-cell exhaustion [37-40], were strongest in T_TOX_ cells at the end of the trajectory (highest pseudotime), whereas the inferred transcription factor activity of *TCF7* was lowest in these cells (Figure 5C). Based on that, we established an signature profile (Supplementary Table 5), to facilitate the identification of terminally exhausted T-cells in scRNA-seq data, for instance using a previously described scoring algorithm [28] (Supplementary Figure 6A).

Plotting the proportion of T_TOX_ cells by sample and pseudotime revealed that terminally exhausted T-cells were most abundant in DLBCL and FL, most variable in MZL, and lowest in MCL (Figure 5D). Clonality analysis based on scTCR data supported this finding by demonstrating that the expansion of PD1^+^ TIM3^+^ T_TOX_ EM_3_ cells was a key feature of the tumor microenvironment of DLBCL, FL and MZL, while MCL and tumor-free LN were predominantly characterized by clonal PD1^-^ T_TOX_ EM_1_ cells (Figure 4A).

### T-cell exhaustion is associated with adverse prognosis in FL and DLBCL

To investigate if T-cell exhaustion is associated with clinical outcome in B-NHL, we extracted a transcriptional signature (see Method section) from terminally exhausted T-cells of our data and applied digital cytometry [41] to bulk RNA data from two large independent retrospective DLBCL cohorts [2, 4]. We found that a higher proportion of terminally exhausted T_TOX_ cells was associated with inferior progression-free survival in both cohorts (Figure 5E, p = 0.003, Figure 5F, p = 0.011). Moreover, the cohort from Schmitz et al. [4] harbored higher proportions of exhausted T-cells in ABC-than GCB-subtype DLBCL (Supplementary Figure 6B), which is in line with a recent flow cytometry-based study [42]. However, there was no difference between ABC- and GCB-subtype DLBCL in our data (Supplementary Figure 4B) or in the cohort from Chapuy et al. [2] (Supplementary Figure 6C). Neither of the genetic subtypes defined in these cohorts were associated with the proportion of exhausted T_TOX_ cells (Supplementary Figure 6B, C). We also evaluated individual somatic mutations (e.g., *MYD88*), amplifications (e.g., *BCL2*), deletions (e.g., 17p), and structural variants (e.g., *BCL6*), which were used to define genetic subtypes by Chapuy et al. [2]. After correction for multiple testing using the Benjamini-Hochberg procedure, none of the genetic aberrations were associated with the proportion of exhausted T_TOX_ cells (Supplementary Figure 6E-H).

We performed a similar analysis using bulk RNA-seq data of a prospective FL cohort of the GALLIUM trial (NCT01332968) [43, 44], and again a higher proportion of exhausted T-cells was associated with inferior survival (Figure 5G, p = 0.04). Overall, these results suggest that T-cell exhaustion is linked to inferior patient outcomes across different B-NHL entities but not clearly associated with cell-of-origin or genetic subtypes.

### IKZF3^+^ T_REG_ EM_2_ are clonally related to T_FH_ cells and associated with grading of FL

In FL and MZL samples, not only T_TOX_ cells but also T_FH_ and T_REG_ EM_2_ cells were clonally expanded (Figure 4A) and significantly enriched (Figure 6A). While T_FH_ cells are well-characterized and known to promote the growth of malignant B-cells in FL [45], the role of LN-derived T_REG_ cells in nodal B-NHL has not been investigated systematically. Aiming to characterize T_REG_ EM_2_ cells, we compared protein and gene expression profiles of T_REG_ EM_2_ and T_REG_ EM_1_ cells, which were evenly distributed across entities. We found that T_REG_ EM_2_ cells were characterized by high protein levels of CD69, ICOS, CD38, PD1, and TIGIT (Figure 6B). At the gene expression level, T_REG_ EM_2_ cells showed high expression of *IKZF3, CXCL13*, and *ASCL2*, but low expression of *KLF2* and *IKZF2* (Figure 6C). We used flow cytometry to confirm the presence of IKZF3 on protein level (Supplementary Figure 7A) and the enrichment of IKZF3^+^ T_REG_ cells in MZL and FL using an independent cohort of 24 LN samples (Supplementary Figure 7B). IKZF2, alias Helios, is well-studied as a marker of natural T_REG_ cells [46], but only few studies have explored the role of IKZF3, alias Aiolos, in T_REG_ cells. A previous study suggested that IKZF3^+^ T_REG_ cells usually lack IKZF2 and represent an inducible rather than natural T_REG_ subset with potent suppressive capacity [47]. Based on this, we intended to identify potential populations related to T_REG_ EM_2_ cells using scTCR data. We found that T_REG_ EM_2_ cells share a substantial proportion of clonotypes with T_FH_ cells (Figure 6D), whereas this was not the case for other T_REG_ cell populations (Supplementary Figure 7C). Based on these aspects, T_REG_ cells could resemble follicular regulatory (Tfr) T-cells, although the latter are reported to express CXCR5 to similar extent as T_FH_ cells [48].

**Figure 6.**
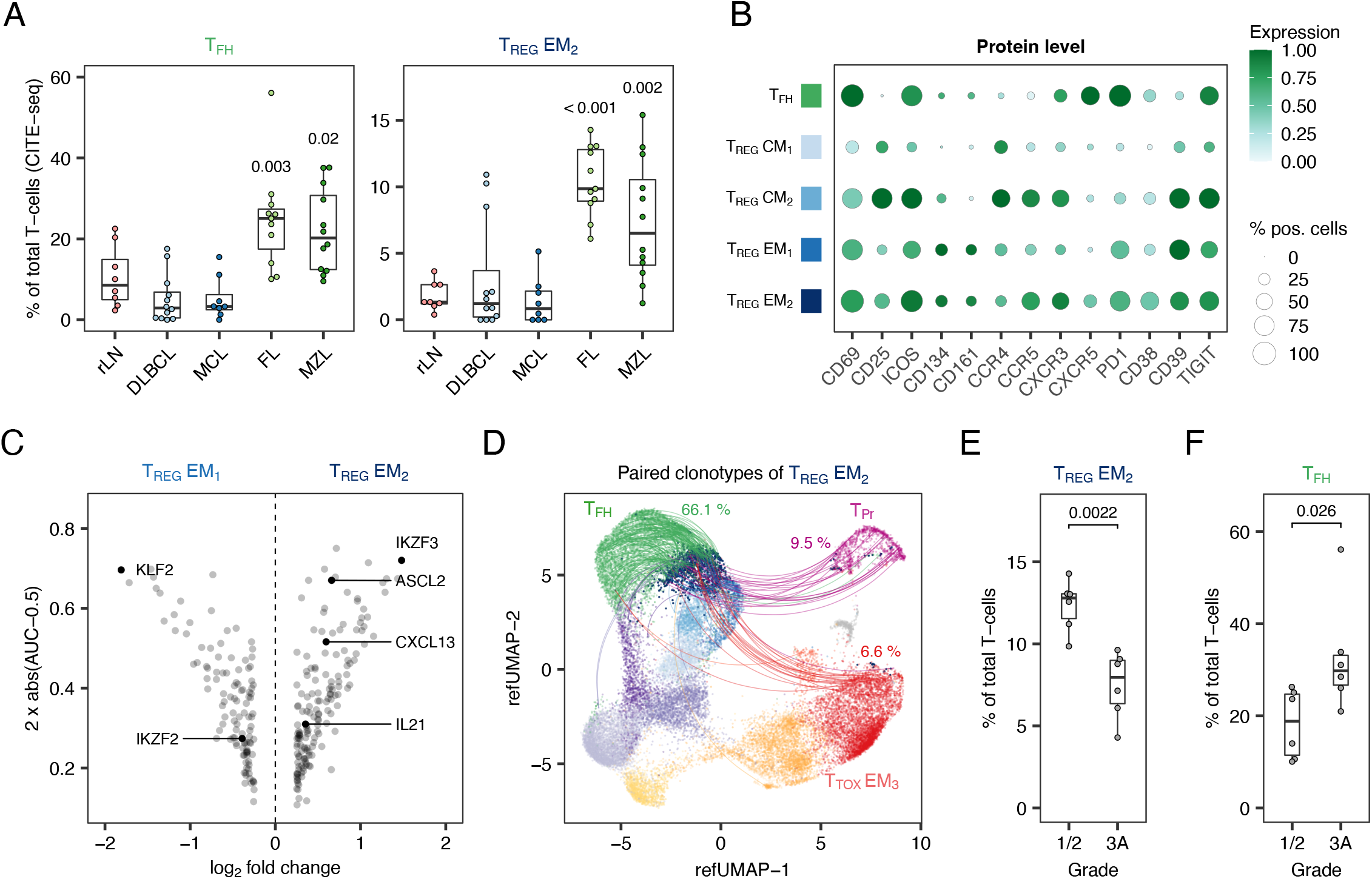
IKZF3+ T_REG_ EM_2_ are clonally related to T_FH_ cells and associated with grading of FL. A) Proportions of TREG EM2 and TFH cells determined by CITE-seq are illustrated as box plots (n = 51). All entities were tested for significance using the two-sided Wilcoxon-test with rLN as reference. B) Dot plot showing the expression of important phenotypic proteins. Size and color of the dots indicate the percentage of positive cells and scaled protein expression, respectively. Values were scaled between 0 and 1. C) Volcano plot illustrating differentially expressed genes between TREG EM2 and EM1 cells. D) 5’ scRNA alongside full-length TCR repertoire data were mapped to the CITE-seq reference data. Lines connect all TREG EM2 cells with any other cell given that both T-cells have the same TCR clonotype. Percentages indicate shares of overlapping clonotypes for TFH, TPr, and TTOX cells. Analysis is based on n = 11 biologically independent patient samples. E-F) Proportions of TREG EM2 (E) and TFH cells (F) determined by CITE-seq are shown in dependence of tumor grading in FL (1/2 versus 3A). Differences were tested for significance using the two-sided Wilcoxon-test.

We further investigated if the proportion of T_REG_ EM_2_ cells was associated with clinical parameters. Lower grade (1/2) FL had a higher proportion of T_REG_ EM_2_ cells (Figure 6E, p = 0.002), whereas higher grade FL (3A) had a higher proportion of T_FH_ cells (Figure 6F, p = 0.03). We applied digital cytometry [41] on the GALLIUM cohort [43, 44] to estimate the proportion of T_REG_ EM_2_ and T_FH_ cells, and to investigate if these subsets were associated with progression-free survival. Indeed, there was a trend towards inferior prognosis in patients with higher proportions of T_FH_ cells (p = 0.05, Supplementary Figure 7D), while no association was found for T_REG_ EM_2_ cells (p = 0.17, Supplementary Figure 7E).

### Highly multiplexed immunofluorescence identifies major T-cell populations within the tumor microenvironment

To localize T-cell subsets in their spatial context and to identify further entity-discriminating features, we used highly multiplexed immunofluorescence in formalin-fixed paraffin-embedded (FFPE) LN tissues using co-detection by indexing (CODEX) [49]. We established a panel of 50 antibodies and two nuclear stains (Supplementary Table 6) and imaged 35 FFPE biopsy cores derived from 19 patient samples (Supplementary Figure 1A, Supplementary Table 1). By clustering marker expression profiles of segmented single cells and validation in high-dimensional fluorescence microscopy images, we identified B-, T-, natural killer (NK), NK T-, mast (MC), plasma (PC), dendritic (DC), follicular dendritic (FDC), and stromal cells, as well as macrophages and granulocytes (Supplementary Figure 8A-C). Among a total of around 5.5 million processed cells, we detected a median of approximately 45,000 T-cells per tissue core (Supplementary Figure 8D), which we further divided into 8 different subpopulations including naïve CD4^+^ and CD8^+^ T-cells (CD45RA^+^), T_FH_ cells (CD45RO^+^, PD1^+^, CXCR5^+^), T_REG_ cells (FoxP3^+^), memory CD4^+^ cells (CD45RO^+^), memory CD8^+^ T_TOX_ cells (CD45RO^+^), exhausted T_TOX_ cells (CD45RO^+^, PD1^+^, TIM3^+^) and T_Pr_ cells (Ki67^+^, Supplementary Figure 8A, C). A high-granularity classification of T-cell subsets, as possible with the CITE-seq and flow cytometry data, was hampered by lower signal-to-noise ratio (e.g., CD62L, CD69) or reduced sensitivity of available antibodies (e.g., IKZF3). To enable a common perspective, we aligned the low-granularity T-cell subpopulations detected by multiplexed immunofluorescence with the 14 high-granularity T-cell subsets identified by CITE-seq (Supplementary Figure 8A). For all 19 LN samples that were analyzed by both approaches, we correlated the proportions for each of the T-cell subsets and observed a median Pearson coefficient of 0.71 across all subpopulations (Supplementary Figure 8E).

### B-NHL disrupts the healthy LN architecture and generates entity-specific microenvironmental patterns

*In-situ* mapping of the above-mentioned cell types and T-cell subpopulations enabled an intuitive visualization of tumor-free or malignant LN structure (Figure 7A, Supplementary Figure 9). To capture spatial organization quantitatively as a means for systematic comparison between tumor-free and malignant LN, we identified the 25 nearest neighbors of each cell by a sliding window approach and tabulated the frequencies of cell types and T-cell subsets in each window. We used k-means clustering on the neighbor frequency tables to identify 10 recurrent neighborhoods (N1-N10, Figure 7B) and PCA to identify the cell types most important for distinguishing N1 to N10 (Figure 7C). This captured important elements of intact LN architecture, including B-cell follicles (N1) with T_FH_- and FDC-rich germinal centers (N2), follicle-surrounding mantle zones (N5), inter-follicular T-cell zones (N6, N7, N9), and sinuses (N10) harboring predominantly stromal cells, macrophages, mast cells, granulocytes, and NK cells (Figure 7A-C). Importantly, this characterization emphasizes the specialized role of T_FH_ cells as they were mostly separated from other T-cell subsets and instead co-localized with FDC and B-cells.

**Figure 7.**
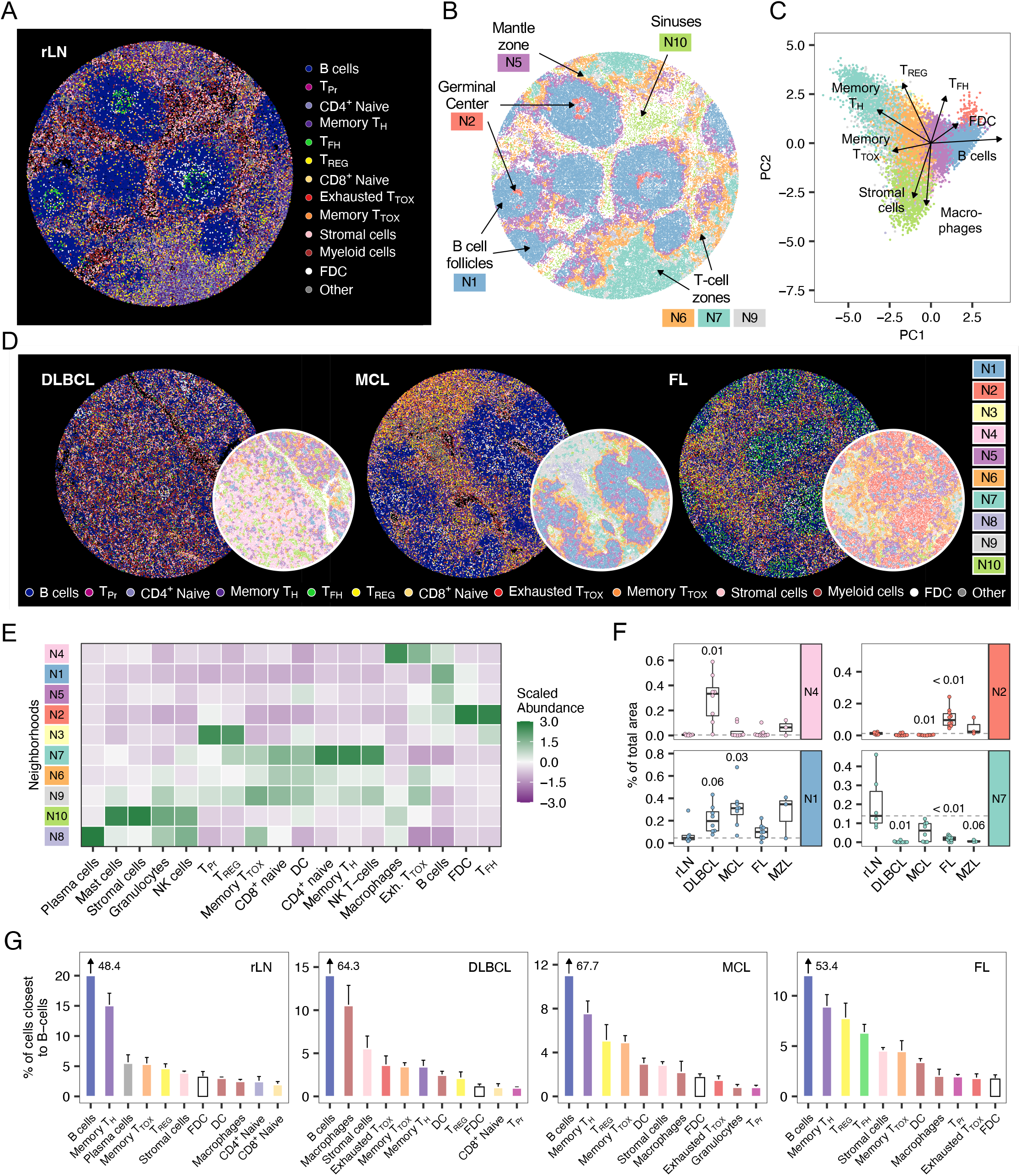
B-NHL disrupts the healthy LN architecture and generates entity-specific microenvironmental patterns. A-B) Each cell of a representative tumor-free LN-derived tissue core (rLN) is colored by its subpopulation (A) or neighborhood (B). C) Neighborhoods shown in panel B were subjected to principal component analysis. The top four loadings of components 1 and 2 are shown as vectors. D) Representative LN-derived tissue cores infiltrated by DLBCL, MCL, or FL. Each cell is colored by subpopulation or neighborhood. E) Heatmap illustrating the mean and column-wise scaled abundance of subpopulations per neighborhood across the complete mIF dataset. F) Box plots showing the proportions of selected neighborhoods of each tissue core (n = 35). Each entity and neighborhood were tested versus rLN using the Wilcoxon-test. P values were corrected for multiple testing using the Benjamini-Hochberg procedure. Dashed lines indicate the median of tumor-free LN (rLN). G) Bar plots showing percentages of cells that were located closest to B-cells. Error bars represent the standard error of the mean. For illustration purposes the B-cell bar is not shown completely but indicated as number. LN: Lymph node.

The described pattern was preserved in all tumor-free tissue cores (Supplementary Figure 10A) but largely disrupted in B-NHL LN (Figure 7D, Supplementary Figure 10B-E). LN-derived tissue cores infiltrated by DLBCL had the least degree of structure and exhibited a diffuse excess of a neighborhood (N4) harboring exhausted T-cells, macrophages and tumor cells, whereas neighborhoods (N7, N8) rich in naïve and memory T_H_ and T_TOX_ cells were absent (Figure 7D-F, Supplementary Figure 10B). Tissue cores from LN infiltrated by FL were characterized by expansion of germinal center-like areas (N2) containing high numbers of (clonal) T_FH_ cells and FDC, surrounded by areas (N6, N9) containing T_REG_, memory T_H_, memory and exhausted T_TOX_, but also B-cells (Figure 7D-F, Supplementary Figure 10D). In tissue cores infiltrated by MCL, we found a significant predominance of follicle-like B-cell areas (N1, Figure 7F), where T_FH_ cells and germinal centers were absent (Figure 7D-E, Supplementary Figure 10C). In contrast to DLBCL and FL, B-cell areas in MCL were well-separated and barely infiltrated by T-cells (Figure 7D, Supplementary Figure 9), resulting in only little contact surface in the transition areas (N6, Figure 7D-E). Entity-specific neighborhood patterns imply different cell-to-cell contacts. To identify interaction partners of B-cells based on spatial proximity, we determined the nearest neighbor of each B-cell and ranked them by frequency (Figure 7G). This analysis suggested strikingly different interaction partner across entities: While B-cells derived from FL, MCL and tumor-free LN were in closest contact to varying T-cell subsets, for instance T_REG_ and T_FH_ cells in FL, we observed that B-cells derived from DLBCL were mostly surrounded by macrophages, stromal and exhausted T-cells (Figure 7G).

In summary, while CITE-seq resolved T-cell heterogeneity quantitatively at a higher granularity, multiplexed immunofluorescence data added a new perspective for studying the LN microenvironment, particularly regarding spatial organization and potential interaction of T-cells with tumor and other immune cells.

## Discussion

Our study provides an in-depth and systematic reference map of LN-derived T-cells in DLBCL, FL, MCL, MZL, and tumor-free samples. Based on more than 100 LN samples, we identified 14 T-cell subsets including their transcriptomic and surface protein signatures, entity-specific abundances, clonality patterns, and a flow cytometry gating strategy. In addition, we employed highly multiplexed immunofluorescence, which resolved T-cell heterogeneity at lower granularity but facilitated us to uncover entity-specific spatial patterns of T-cell infiltration.

Across all entities, malignant LN were characterized by loss of CD4^+^ and CD8^+^ naive T-cells, as well as CD69^-^ T_H_ CM_1_ and CD69^+^ CM_2_ cells, but to variable extents harbored clonally expanded CD4^+^ and CD8^+^ PD1^+^ TIM3^+^ T_TOX_ cells. The latter were part of a cellular trajectory that continuously converged into terminally exhausted T-cells. Beside high expression of various inhibitory receptors (PD1, TIM3, LAG3, TIGIT, CD39) and reduced expression of effector molecules (*GZMA, GZMH*), these cells had lost *TCF7* transcription factor activity, which is considered a crucial and early event towards terminal T-cell exhaustion [50]. Higher proportions of these cells were not significantly linked to cell-of-origin or genetic subtypes in DLBCL but instead associated with inferior survival in FL and DLBCL patients. These results are in line with previous studies that used TIM3 or LAG3 as surrogate markers to estimate T-cell exhaustion in DLBCL [51] or FL patients [52]. PD1^+^ TIM3^+^ T_TOX_ cells were located within several spatial neighborhoods, thereby allowing for contact with various types of immune cells. Specifically in LN infiltrated by DLBCL, PD1^+^ TIM3^+^ T_TOX_ cells were strongly co-localized with tumor cells and macrophages, which have recently been shown to be attracted by exhausted T-cells and inversely, to reinforce T-cell exhaustion [53]. This observation might explain why DLBCL harbored the highest proportions of PD1^+^ TIM3^+^ T_TOX_ and why higher numbers of macrophages are associated with inferior outcome in DLBCL patients [54, 55]. While DLBCL was also characterized by the least degree of structure, we observed that LN infiltrated by MCL exhibited a rather strict separation of tumor and T-cell areas, resulting in less possibility for direct cellular interaction. As ongoing antigen presence is a prerequisite of T-cell exhaustion [50], the remarkably low proportions of PD1^+^ TIM3^+^ T_TOX_ cells might be the consequence of such maintained compartmentation. Further effort is needed to investigate how these distinct spatial patterns are formed and how specific cellular interactions, for instance with macrophages, may impact on re-directed T-cells.

While the majority of clonal T_TOX_ cells in B-NHL could be assigned to the TCF7^-^ trajectory, we identified a second trajectory that was characterized by expression of PD1, absence of TIM3 and other inhibitory receptors, but maintained transcription factor activity of *TCF7*. Previous work suggested that both subsets play opposite roles, as, for instance, only TCF7^+^ PD1^+^ T-cells can be reinvigorated upon checkpoint inhibition, whereas terminally exhausted TCF7^-^ T-cells cannot be restored [56, 57]. This observation is of particular interest because immune-checkpoint blockade is remarkably ineffective in nodal B-NHL [58, 59], which might be – in the light of our study – due to the predominance of terminally exhausted T-cells. Indeed, also Zheng et al. found that B-NHL seem to harbor higher levels of terminally exhausted T-cells than most other cancer entities [60]. However, we speculate that B-NHL patients harboring high levels of PD1^+^ TIM3^-^ T_TOX_ cells could represent a subgroup that benefits most from combining T-cell engaging immunotherapies and immune checkpoint blockade.

Beyond clonal T_TOX_ cells, we found that both FL and MZL were characterized by clonal expansion of T_FH_ and IKZF3^+^ T_REG_ EM_2_ cells, and overall had a similar pattern of T-cell infiltration. Whereas T_FH_ cells have been extensively studied and are known to support the growth of malignant B-cells in FL [45], an enrichment of IKZF3^+^ T_REG_ EM_2_ cells has, to our knowledge, neither been described in FL nor MZL. Even recent scRNA-seq studies of FL have not captured such a subset, possibly because the heterogeneity of T_REG_ cells was not investigated in detail or because FL was not compared to other B-NHL entities [18, 61]. We observed higher proportions of IKZF3^+^ T_REG_ EM_2_ cells in low-graded FL patients, suggesting that this T-cell subset could modulate the proliferation capacity of malignant B-cells. IKZF3^+^ T_REG_ EM_2_ cells are suggested to bear strong suppressive capacity and represent an induced T_REG_ phenotype [47]. Indeed, we demonstrated that IKZF3^+^ T_REG_ EM_2_ cells and T_FH_ cells carry a substantial proportion of identical TCR clonotypes, which implies that IKZF3^+^ T_REG_ EM_2_ cells most likely derive from T_FH_ cells. The close relation of both subsets is further substantiated by the fact that, on the one hand, T_FH_ cells express IKZF3 constitutively [62] and, on the other hand, IKZF3^+^ T_REG_ EM_2_ cells express high levels of PD1, *CXCL13* and *ASCL2 –* known marker genes and proteins of T_FH_ cells [27, 63]. The induction of FoxP3, or more generally, the conversion of T (follicular) helper cells into T_REG_ cells can be mediated by augmented stimulation or auto-inflammatory conditions and has been previously described for B-NHL derived T-cells [64-66]. More mechanistically oriented studied are needed to clarify whether IKZF3^+^ T_REG_ indeed derive from T_FH_ cells and whether they can be leveraged to beneficially affect the course of disease.

While highly multiplexed immunofluorescence provides insights that cannot be obtained by suspension-based single cell assays [67], its application in larger patient cohorts and achieving a more fine-grained resolution of certain cell subsets are still challenging. Particularly in FFPE tissue, which is routinely used for diagnostic samples, development of antibody panels requires labor-intensive optimization and validation, and certain markers may not be detectable due to loss of antigenic reactivity. More importantly, current computational approaches represent a bottleneck, as significant manual operation is needed to annotate and validate cell types, which is particularly relevant when handling large tissue areas with millions of cells. Further improvements of computational approaches will enable a broader application of highly multiplexed immunofluorescence across large clinical cohorts [68-70].

In summary, our work refines previous knowledge of lymphoma-infiltrating T-cells by employing recent technological advances and offers a new perspective on different B-NHL entities, which have previously not been studied. This broader yet more detailed view revealed that different B-NHL entities shape their T-cell microenvironment in distinct manners, which could not be readily detected in studies investigating only single entities. We generated a multimodal resource that facilitated deciphering both beneficial and adverse directions of T-cell response in nodal B-NHL and will thus contribute to improving T-cell engaging treatment approaches.

## Material and Methods

### Lymph node samples

Our study was approved by the Ethics Committee of the University of Heidelberg (S-254/2016). Informed consent of all patients was obtained in advance. LN patient samples were processed and frozen until further analysis as previously described [71]. Samples from patients after allogeneic stem cell transplantation, CAR T-cell or bispecific antibody therapy were not used in this study to minimize treatment-associated effects on the T-cell microenvironment. For the same reason, samples were collected earliest 3 months after cessation of the last treatment.

### Single-cell 3’ RNA-seq and epitope expression profiling

Cells were thawed and immediately washed to remove DMSO. In order to prevent entity-associated batch effects, samples were processed in batches of four to five containing at least three different entities. After thawing, we applied a dead cell removal kit (Miltenyi Biotec) to all samples to reach a viability of at least 85 to 90 %. Samples not reaching a viability above 85 % were excluded. Then, 5 × 10^5^ viable cells were stained by a pre-mixed cocktail of oligonucleotide-conjugated antibodies (Supplementary Table 2) and incubated at 4 °C for 30 minutes. Cells were washed three times with ice-cold washing buffer and each time centrifuged at 4 °C for 5 minutes. After completion, cells were counted and viability was determined again. Samples not reaching a viability above 85 % were excluded. The preparation of the bead-cell suspensions, synthesis of complementary DNA and single-cell gene expression and antibody-derived tag (ADT) libraries were performed using a Chromium single-cell v3.1 3[kit (10x Genomics) according to the manufacturer’s instructions.

### Single-cell 5’ RNA-seq and T-cell receptor repertoire profiling

Apart from epitope staining, sample processing was identical to 3’ scRNA-seq. The preparation of the bead-cell suspensions, synthesis of complementary DNA and single-cell gene expression and TCR libraries were performed using a Chromium single-cell v2 5[and human TCR amplification kit (both 10x Genomics) according to the manufacturer’s instructions.

### Single-cell library sequencing and data processing

3’ gene expression and ADT libraries were pooled in a ratio of 3:1 aiming for 40,000 reads (gene expression) and 15,000 reads per cell (ADT), respectively. Sequencing was performed on a NextSeq 500 (Illumina). 5’ gene expression libraries were sequenced on a NextSeq 2000 (Illumina) aiming for 50,000 reads per cell. TCR libraries were sequenced on a NextSeq 500 (Illumina) aiming for a minimum of 5,000 reads per cell. After sequencing, the Cell Ranger (10x Genomics, v6.1.1) function *cellranger mkfastq* was used to demultiplex and to align raw base-call files to the reference genome (hg38). For 3’ gene and epitope expression libraries, the obtained FASTQ files were counted by the *cellranger count* command, whereas *cellranger multi* was used for 5’ gene expression and TCR libraries. As reference for TCR libraries, the VDJ ensembl reference (hg38, v5.0.0) was used. If not otherwise indicated default settings were used for all functions.

### Analysis and integration of CITE-seq data

The R package Seurat (v4.1.0) was used to perform data quality control, filtering, and normalization. Gene counts per cell, ADT counts per cell and percentages of mitochondrial reads were computed using the built-in functions. Principal component analysis, shared nearest neighbor (SNN) based clustering and UMAP were done based on the combined transcriptome and epitope data. After mapping the CD3 and CD19 epitope expression, non-T-cell clusters and doublets were removed. For data integration across the different preparation batches, we used the IntegrateData function of the Seurat package. For multimodal clustering based on gene and epitope expression, the weighted nearest neighbor approach was used [22]. To estimate the proportion of positive cells for each surface marker, we calculated the denoised protein expression using the totalVI python package [72].

### Inferring transcription factor activity based on single-cell gene expression

We used the SCENIC (Single-Cell rEgulatory Network Inference and Clustering) package [23] to infer gene regulatory networks and transcription factor activity based on scRNA-seq data. Functions were used according to publicly available vignettes.

### Surface and intracellular flow cytometry staining

LN-derived cells were thawed and stained for viability using a fixable viability dye e506 (Thermo Fisher Scientific) and for different surface markers depending on the experimental set-up. The following surface antibodies were used: anti-CD3-PerCP-Cy5.5, anti-CD4-PE-Dazzle, anti-CD8-APC-Cy7, anti-CD45RA-FITC, anti-CD25-BV421, anti-CD31-BV605, anti-CXCR5-BV711, anti-TIM3-BV711, anti-CD278-BV605, anti-PD1-PE-Cy7, anti-CD69-AF700, anti-CD244-BV421 (all BioLegend). For subsequent intracellular staining, cells were fixed and permeabilized with the intracellular fixation/permeabilization buffer set (Thermo Fisher Scientific) and stained with anti-Ki67-BV785, anti-FoxP3-AF647, anti-IKZF3-PE or adequate isotype controls (Thermo Fisher Scientific, BD Biosciences). Then, cells were analyzed using an LSR Fortessa (BD Biosciences) and FACSDiva (BD Biosciences, version 8). For analysis and gating of flow cytometry data FlowJo (v10.8.0) was used.

### Multinomial classification of T-cell subpopulations

First, we evaluated whether multimodally defined T-cell subsets can be distinguished in general by using surface markers only. Therefore, we trained gradient boosting models (“xgbTree”) [29] on the basis of surface marker expression of single-cell data. To reduce data load, only 30 % of all cells were used. 10-fold cross-validation was employed to optimize the model. Since surface marker were not sufficient to reach sufficient accuracy, additional models were trained using surface marker plus gene expression of MKI67, IKZF3, and FoxP3, since these genes were differentially expressed between T-cell subsets that could not be sufficiently predicted using only surface proteins. Finally, markers were ranked and selected by their variable importance to build flow cytometry panels. In case two or more markers deliver similar information (e.g., CD95 and CD45RA), only one was selected. Gating strategies were built using the R package hypergate [30] and optimized in an iterative process. The final gating strategy for all multimodally defined T-cell subsets is illustrated in Supplementary Figure 2C and 3.

### Prediction of B-NHL entity from T-cell proportions

To assess feasibility of predicting B-NHL entity or tumor-free condition based on the proportions of all T-cell subsets and overall T-cell frequency, we trained classifiers based on multivariate regression with L1 penalty (LASSO) implemented in the R package glmnet (v4.1) [73]. The hyperparameter lambda was determined using cv.glmnet with balanced folds and weights inversely proportional to class size. The confusion matrix was computed using leave-one-out cross-validation.

### Analysis of T-cell receptor diversity

To estimate the diversity of the TCR repertoire based on single-cell TCR profiling, we employed the R package immunarch (v0.8.0) [74] on the output files of the cellranger pipeline. TCR diversity across samples was compared using a rarefraction analysis. Therefore, the function repDiversity with method = “raref” was applied.

### Mapping of 5’ scRNA-seq data onto CITE-seq reference data

To evaluate scTCR data in the context of multimodally defined T-cell subsets, 5’ scRNA-seq data were mapped onto the CITE-seq reference data, using the built-in functions *FindTransferAnchors* and *MapQuery* of the Seurat package (v4.1.0). The mapping accuracy was evaluated by comparing the T-cell subset proportion of 5’ and 3’ data of the identical patient sample.

### Pseudotime analysis and exhaustion signature

Pseudotime analysis based on gene expression profiles of T_TOX_ cells was performed using the monocle3 package [75, 76]. Briefly, the Seurat object was converted into a cell data set. Trajectory and pseudotime analysis were performed using the functions learn_graph and order_cells, respectively. As root cells, naive CD8^+^ T-cells were selected. Minimal branch length was set to 10, otherwise default settings were used.

To define a transcriptional module for T-cell exhaustion, differentially expressed genes of terminally exhausted T-cells, meaning T_TOX_ cells with highest levels of inhibitory receptors and decreasing expression of effector molecules (Figure 5C), were determined (Supplementary Table 5). The UCell package (v1.3.1) [28], which is based on the Mann-Whitney U statistic, was then applied to calculate an exhaustion score for each cell.

### Deconvolution of bulk RNA sequencing data

Deconvolution of bulk RNA-seq data was performed using the interactive web application (https://cibersortx.stanford.edu/) developed by Newman and colleagues [41]. First, a signature matrix was created based on the scRNA-seq data and the cluster annotation as cell types.

Minimum expression was set to zero and 200 replicates were used. Otherwise, default settings were applied to create a signature matrix. Second, cell fractions were imputed using bulk RNA-seq data and signature matrix as input. S-mode batch correction and absolute mode was used for each analysis.

### Survival analysis

Survival data were only obtained from previously published data or studies [2, 4, 43]. Analysis of the progression-free survival probability was performed in combination with the estimated cell type proportions using deconvolution of bulk RNA-seq data. To divide the data into two groups (low, high), we determined a cut-off based on the maximized p value of a log-rank test using the maxstat R package (v0.7.25). Kaplan-Meier curves were drawn using the survminer R package (v0.4.9).

### Tissue microarray and coverslip preparation

Representative tumor or tumor-free lymph node areas in archival FFPE tissue blocks from 19 patients (Supplementary Figure 1A, Supplementary Table 1) were selected by board-certified pathologists of the Tissue Bank of the National Center for Tumor Diseases and Institute of Pathology at the University Hospital Heidelberg. Tissue microarrays (TMA) containing two 4.5 mm cores per patient were generated. TMA sections (4 μm) were mounted onto Vectabond-precoated 25 × 25 mm coverslips and coated in paraffin for storage until staining [77].

### Antibody conjugation, validation and titration

Multicolor immunofluorescence was performed using the co-detection by indexing (CODEX) approach [49]. Antibodies used for CODEX experiments are summarized in Supplementary Table 6. Purified, carrier-free antibodies (50-100 μg per reaction) were reduced with Tris(2-carboxyethyl)phosphine (TCEP) and conjugated at 1:2 weight/weight ratio to maleimide-modified CODEX DNA oligonucleotides, which were purchased from TriLink Biotechnologies and deprotected via retro-Diels-Alder reaction. Conjugated antibodies were first evaluated in CODEX singleplex stains on tonsil and/or lymphoma tissue by comparison with online databases (The Human Protein Atlas, Pathology Outlines), immunohistochemical reference stains and/or published literature under the supervision of a board-certified pathologist. Staining patterns were validated in multiplex experiments in the presence of positive and negative control antibodies and the appropriate dilution of each antibody was titrated starting from 1:100 to optimize signal-to-noise ratio.

### Multiplex tissue staining and fixation

Coverslips were deparaffinized, rehydrated and submitted to heat-induced epitope retrieval at pH9 (Dako target retrieval solution, #S236784-2, Agilent) and 97°C for 10 min in a Lab Vision PT module (Thermo Fisher). After blocking of non-specific binding with CODEX FFPE blocking solution, coverslips were stained overnight with the full antibody panel at the dilutions given in Supplementary Table 6 in CODEX FFPE blocking solution [78] in a sealed humidity chamber at 4 °C on a shaker. Coverslips were then fixed with 1.6% paraformaldehyde, followed by methanol and BS3 fixative (Thermo Fisher) before storage in CODEX buffer S4 until imaging [78].

### Multicycle Imaging

Stained coverslips were mounted onto custom acrylic plates (Pololou Corporation) with mounting gaskets (Qintay), thereby creating a flow cell with a surface area of 19 × 19 mm above the tissue for fluid exchange. Acrylic plates were inserted into a Keyence BZ-X710 inverted fluorescence microscope equipped with a CFI Plan Apo λ 20x/0.75 objective (Nikon) using custom adapters. For each core an area of 7×7 fields of view (30% overlap between tiles) and an adequate number of z planes (10-14) required to capture the best focal plane across the imaging area were selected. Multicycle imaging was performed using a CODEX microfluidics device and CODEX driver software v1.29.0.1 (Akoya Biosciences). Exposure times and assignment of markers to imaging cycles and fluorescent channels are provided in Supplementary Table 6. After completion of multicycle imaging, coverslips were stained with hematoxylin/eosin and the same areas were imaged in brightfield mode.

### Image processing

Raw TIFF images were processed using the RAPID pipeline [79] in Matlab (version R2020a) with the following settings: nCyc=51, nReg= number of regions imaged (depending on TMA), nTil=49, nZ= number of z-planes imaged (depending on TMA), nCh=[1,4], nTilRow=7, nTilCol=7, overlapRatio=0.3, reg_range=1:nReg, cyc_range=1:nCyc, til_range=1:nTil, cpu_num=depending on computer system used, neg_flag=1, gpu_id=depending on number of GPUs available, cyc_bg=1. After deconvolution (two iterations), best focal plane selection, lateral drift compensation, stitching of individual images and background subtraction, processed images were concatenated to hyperstacks. All tissue cores were checked visually for staining quality of each antibody in ImageJ/Fiji (version 1.53q).

### Cell segmentation and cell type annotation

Individual nuclei were segmented based on the Hoechst stain (cycle 1), derived nuclear masks were dilated, and cellular marker expression levels were quantified using a modified version of the Mask region-convolutional neural network (R-CNN) based CellSeg software [80] run on the full resolution RAPID stitched images with the following parameters: GROWTH_PIXELS_PLANE = 1.0, output_adjacency_quant = True, BOOST = 1, OVERLAP = 80, MIN_AREA = 80, INCREASE_FACTOR = 4.0, AUTOBOOST_PERCENTILE = 99.98. A threshold based on the intensity of the nuclear markers Hoechst and DRAQ5 was used to exclude non-cellular events and remove cells from tissue areas of low image quality. Marker expression levels compensated for lateral spillover were used for further analysis and the range of each marker was z normalized per imaging run. A total of n=5,690,284 cells were submitted to an initial round of Leiden-based clustering (n_neighbors=10, resolution=2) on key phenotypic markers (CD11b, CD11c, CD14, CD15, CD16, CD163, CD20, CD206, CD21, CD25, CD3, CD31, CD34, CD38, CD4, CD45, CD5, CD56, CD57, CD68, CD7, CD79a, CD8, CD90, FOXP3, HLA-DR, kappa light chain, lambda light chain, MCT, PAX5, PDPN) using the scanpy Python package [81], as previously described [82]. Each of the resulting clusters (n = 55) was assessed for purity of its cell type composition based on marker expression and overlays of the cells in each cluster onto image hyperstacks using CODEX scripts for ImageJ/Fiji (available at https://github.com/bmyury/CODEX-fiji-scripts). Clusters were merged, split, and/or further subclustered as appropriate to define broad cell types (e.g., T-cells, B cells, etc.). This process was repeated within these cell types using additional markers as appropriate to annotate more granular cell subsets. Briefly, marker combinations used for cell typing of non-T-cells include CD16, CD68, CD163, CD206 and HLA-DR (macrophages); CD11c, CD68 and HLA-DR (DC); CD15 (granulocytes); MCT and GRZB (mast cells); CD34, CD31, CD90 and PDPN (stromal cells); PDPN and CD21 (FDC); CD56 (NK and NKT-cells); CD38, CD31, and kappa and lambda light chain (plasma cells); PAX5, CD20 and CD79a (B-cells). Further, T-cell subsets were derived based on key subset markers (CD45, CD45RA, CD45RO, CD3, CD5, CD7, CD4, CD8, FOXP3, CXCR5, CXCL13, PD1, TIM3, CD31, and Ki67) and were validated using expression of additional T-cell markers in the panel and overlays onto image hyperstacks.

### Neighborhood and nearest neighbor analysis

Neighborhood analysis was modified based on a previously described approach [49]. For each cell of the joint highly multiplexed immunofluorescence data, the 25 nearest neighbors were determined based on their Euclidean distance of the X and Y coordinates, resulting in one ‘window’ of cells per individual cell. Next, these windows were grouped using k-means clustering based on the cell type proportions within each window. Finally, each cell was annotated by the neighborhood of its surrounding ‘window’. K = 10 was selected based on the overlays of the neighborhood assignments with the original fluorescent and H&E-stained images. Higher values of k did not result in an improved biologically interpretable number of neighborhoods.

### Interactive browsing of highly multiplexed immunofluorescence data

All tissue cores imaged in this study including staining of 52 different markers are available for interactive browsing at http://45.88.80.128:8765.

### Data availability

All single-cell gene expression, epitope and TCR data will be available in the HeiData database (https://heidata.uni-heidelberg.de) under accession number 0SNSFB upon publication. Highly multiplexed immunofluorescence images are available in the BioStudies database (https://www.ebi.ac.uk/biostudies/) under accession number S-BIAD565 [83] upon publication.

### Code availability

The computational codes, in the form of Rmarkdown documents, for reproducing all main and supplementary figures will be available at github.com upon publication.

## Acknowledgements

T.R. was supported by a fellowship of the German Federal Ministry of Education and Research (BMBF) and a physician scientist fellowship of the Medical Faculty of University Heidelberg. M.A.B was supported by a Career Development award of the International Myeloma Society (IMS). D.F. was supported by the PhD program of the European Molecular Biology Laboratory (EMBL). H.V. was supported by a fellowship of the German Federal Ministry of Education and Research (BMBF). N.L. was supported by a Heidelberg School of Oncology (HSO2) fellowship from the National Center for Tumor Diseases (NCT) Heidelberg. O.W. is supported by an Else-Kröner Excellence Fellowship from the Else-Kröner-Fresenius Stiftung (Project-ID 2021_EKES.13). M.S. was supported by a grant of the Deutsche Forschungsgemeinschaft (DFG). S.D. was supported by a grant of the Hairy Cell Leukemia Foundation, the Heidelberg Research Centre for Molecular Medicine (HRCMM) and an e:med junior group grant of the German Federal Ministry of Education and Research (BMBF). For the data management we thank the Scientific Data Storage Heidelberg (SDS@hd) which is funded by the state of Baden-WuCrttemberg and a DFG grant (INST 35/1314-1 FUGG). We thank Carolin Kolb (University Hospital Heidelberg) and the EMBL flow core facility, and the DKFZ Single-Cell Open Lab (scOpenLab) for their excellent (technical) assistance. The GALLIUM study (NCT01332968) was sponsored by F. Hoffmann-La Roche.

## Author Contributions

Conceptualization, T.R., M.A.B., W.H., S.D.; Software, H.V.; Validation T.R., M.A.B., L.L-C., C.M.S., M.S.; Formal Analysis T.R., M.A.B., D.F., H.V., F.C., V.P., W.H.; Investigation, T.R., M.A.B., D.F., M.K.; Resources, N.L., A.B., G.M., C.M., O.W., G.P.N., W.H., S.D.; Writing – Original Draft, T.R., M.A.B., B.J.B., W.H., S.D.; Writing – Review & Editing, all; Visualization, T.R., W.H.; Supervision, T.R., C.M.S., G.P.N., W.H., S.D.; Project Administration, T.R., W.H., S.D.; Funding Acquisition, G.P.N., W.H., S.D.

