## Supplementary Figures for "Multimodal and spatially resolved profiling identifies distinct patterns of T-cell infiltration in nodal B-cell lymphoma entities"

### Supplementary Figure 1

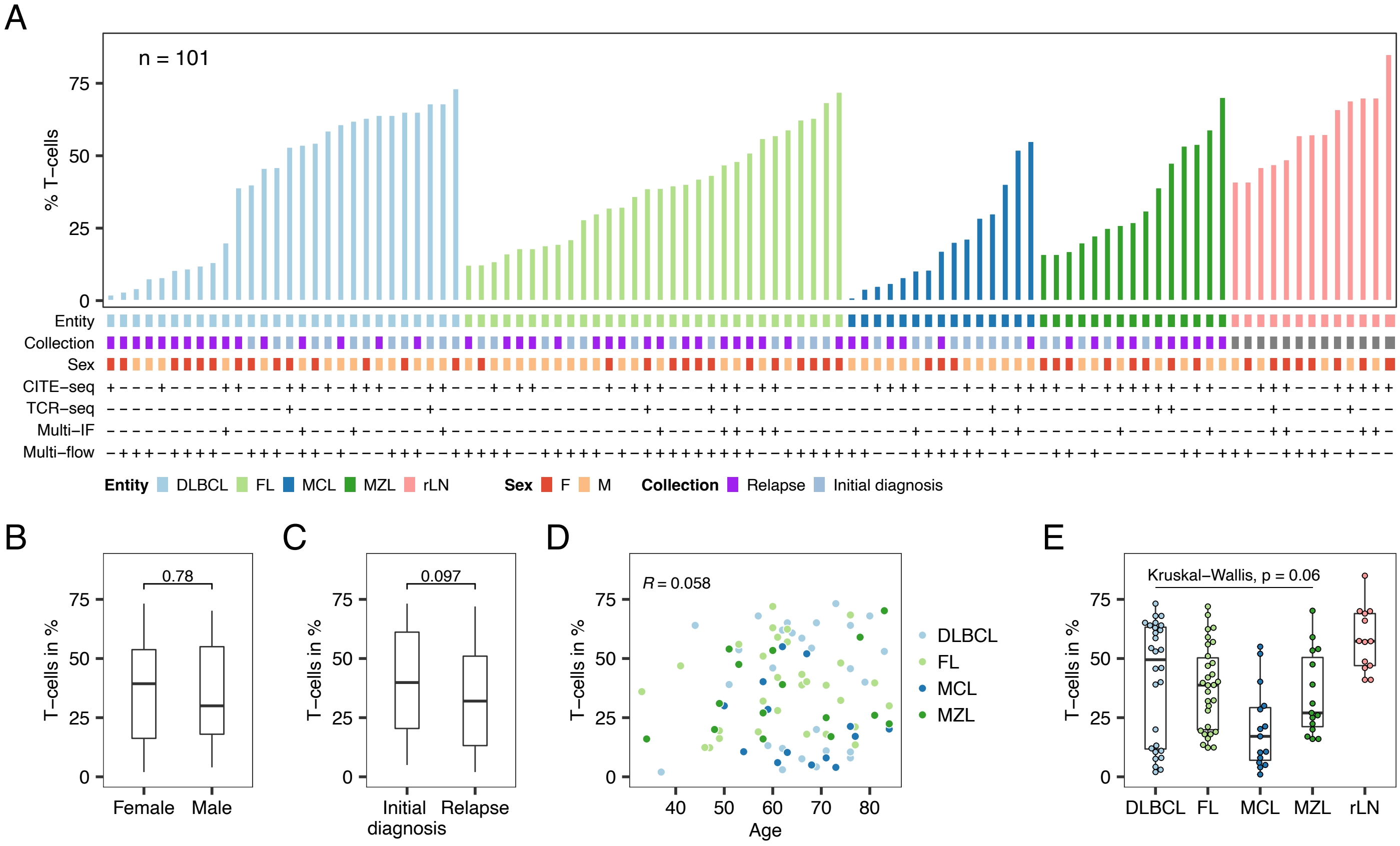


**Supplementary Figure 1.**

A) Overview showing entity, pre-treatment status, sex, assay availability and the overall T-cell proportion of a total 101 LN samples used in this study. B-E) Box or scatter plots illustrating the associations of patient characteristics with the overall T-cell proportion determined by flow cytometry. P values and/or correlation coefficients were calculated using the Wilcoxon-test (B, C), Pearson's linear correlation (D) or Kruskal-Wallis-test (E). LN: Lymph node.

### Supplementary Figure 2

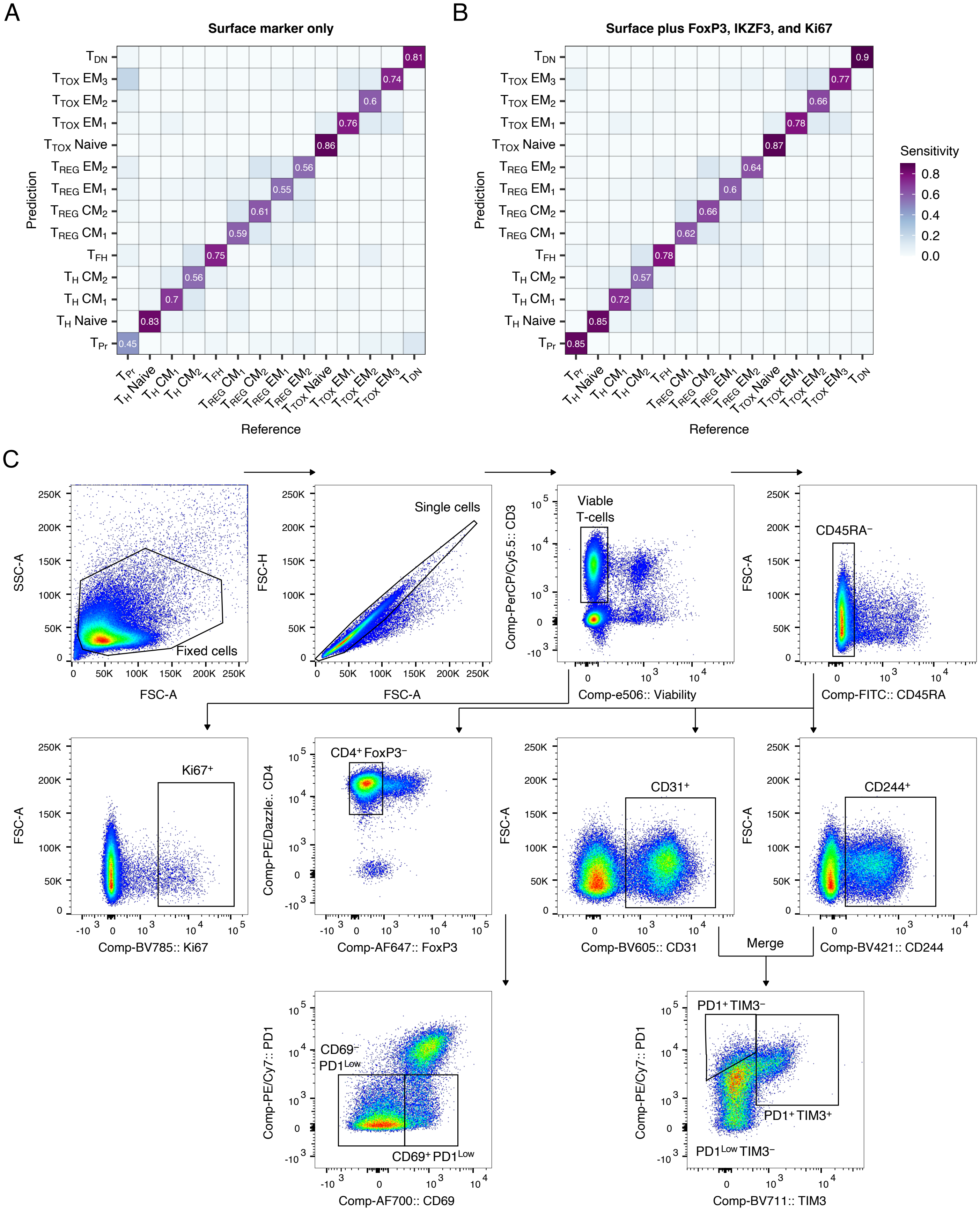


**Supplementary Figure 2.**

A-B) Gradient boosting classifiers were trained and tested to evaluate whether T-cell subsets identified by CITE-seq can be predicted on basis of surface markers (A) or surface markers and differentially expressed intracellular markers accessible by flow cytometry (B, FoxP3, IKZF3, Ki67). Shown is the proportion of correctly predicted cells per T-cell subset. C) Flow cytometry gating strategy for one representative lymph node sample (see method section for details) for $T_{Pr}$ (Ki67⁺), $T_H$ $CM_1$ (CD69⁻ PD1^Low), $T_H$ $CM_2$ (CD69⁺ PD1^Low), $T_{TOX}$ $EM_1$ (PD1^Low TIM3⁻), $T_{TOX}$ $EM_2$ (PD1⁺ TIM3⁻), and $T_{TOX}$ $EM_3$ (PD1⁺ TIM3⁺) cells.

### Supplementary Figure 3

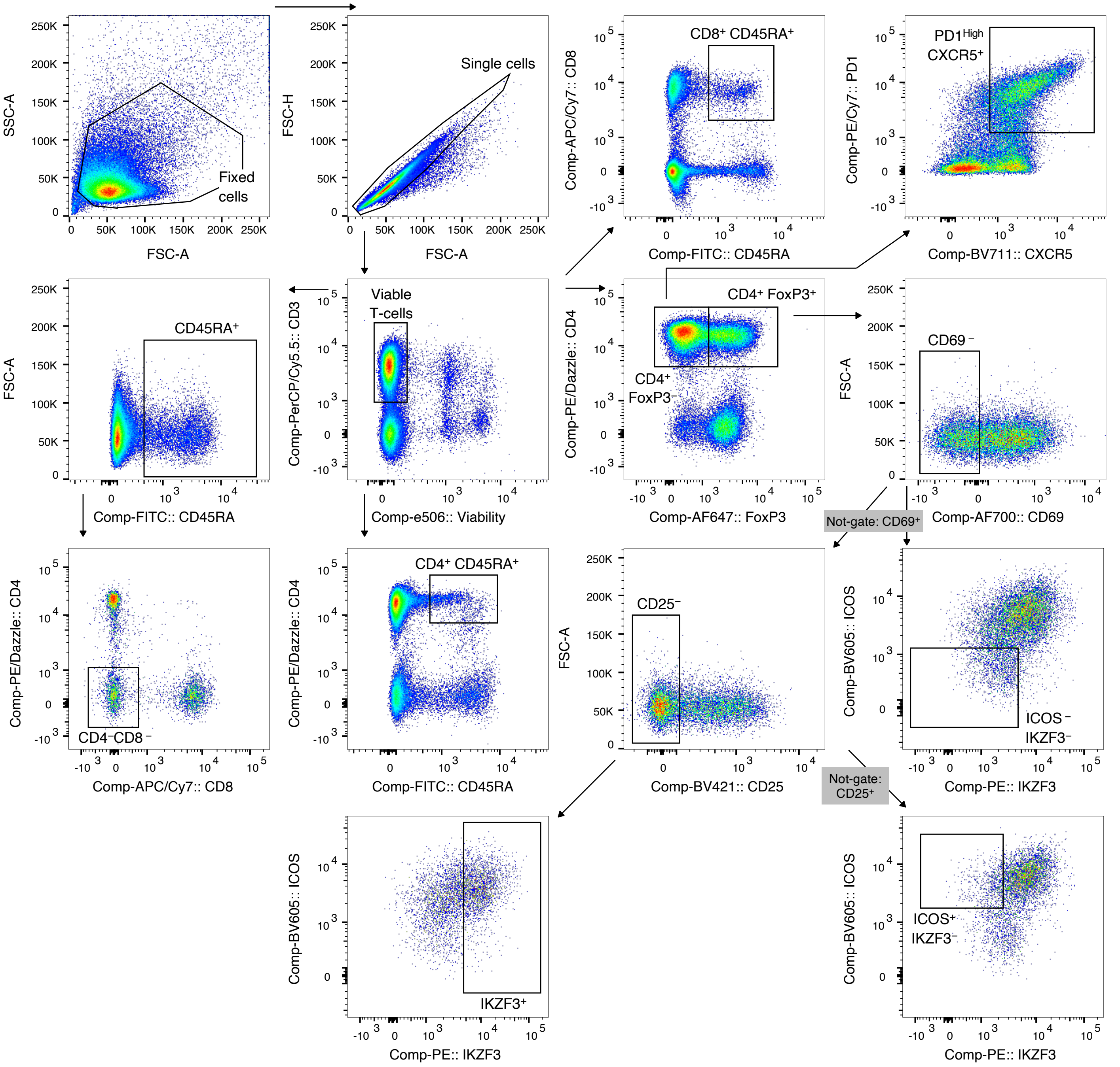


**Supplementary Figure 3.**
Flow cytometry gating strategy (see method section for details) for one representative LN sample for $T_H$ Naïve (CD4+ CD45RA+), $T_{TOX}$ Naïve (CD8+ CD45RA+), $T_{DN}$ (CD4− CD8−), $T_{FH}$ (PD1$^{High}$ CXCR5+), $T_{REG}$ $CM_1$ (CD69−), $T_{REG}$ $CM_2$ (ICOS+ IKZF3−), $T_{REG}$ $EM_1$ (ICOS− IKZF3−), and $T_{REG}$ $EM_2$ (IKZF3+). LN: Lymph node.

### Supplementary Figure 4

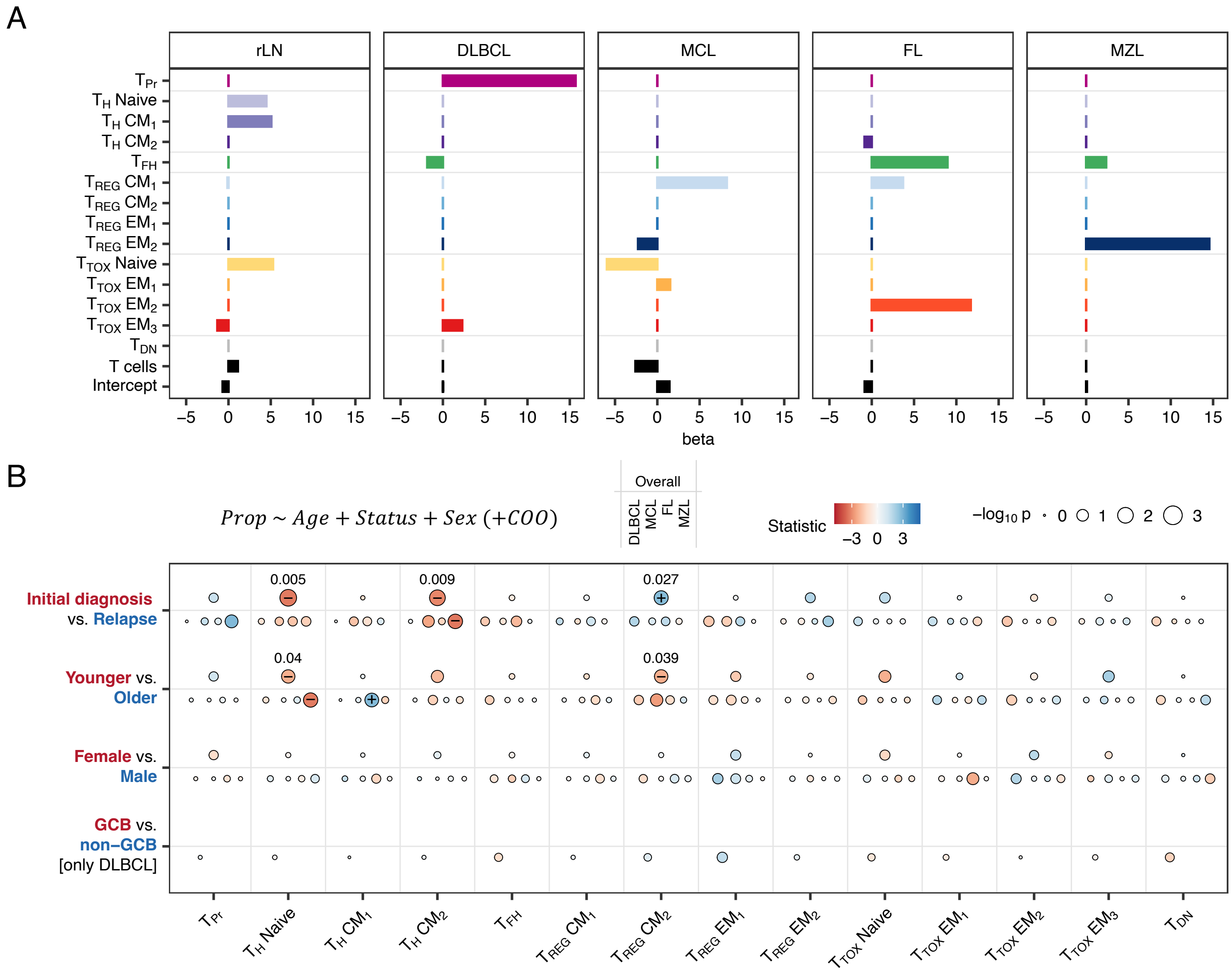


**Supplementary Figure 4.**

A) Beta coefficients derived from a LASSO-regularized multinomial regression model predicting rLN, MZL, FL, MCL, or DLBCL based on the subset and overall T-cell proportions. B) Multivariate linear models were fit using sex, age, treatment status, COO (only DLBCL) as covariates, and the proportion of each T-cell subset as dependent variable. Models were fit for the total dataset and for each entity separately, as indicated. Diameter and color of the dots indicate –log10 p value and size of the coefficient, respectively. P values were corrected using the Benjamini-Hochberg procedure and significant p values are indicated by +/–. COO: Cell-of-origin.

### Supplementary Figure 5

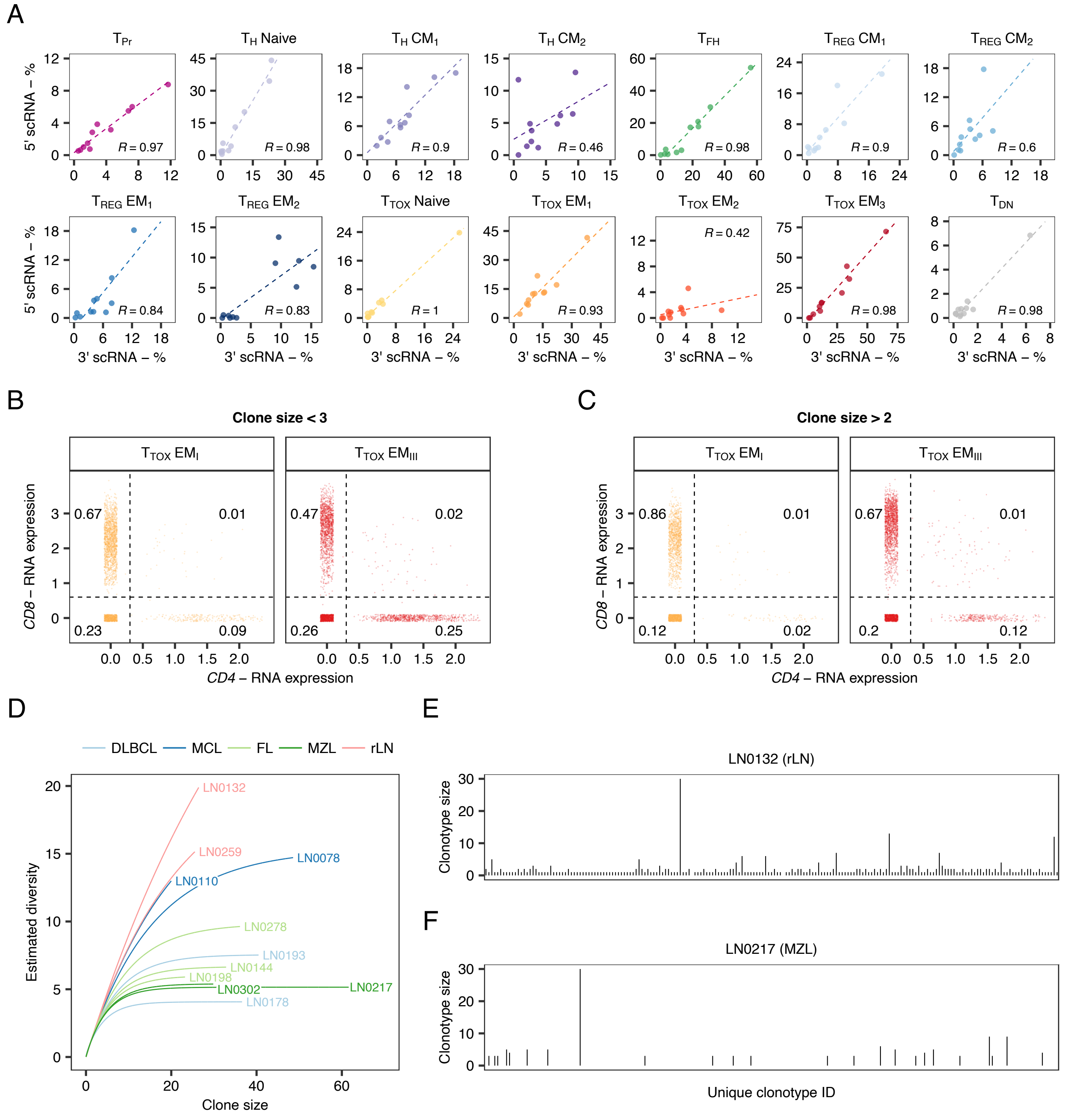


**Supplementary Figure 5.**

5' scRNA alongside full-length TCR repertoire data from n = 11 biologically independent LN patient samples were mapped to the CITE-seq reference data. A) T-cell subset proportions determined by 5' scRNA or CITE-seq reference data were correlated across 11 patient samples and 14 multimodally defined T-cell subsets. R values represent Pearson's linear correlation coefficients. B-C) CD4 and CD8 gene expression of $T_{TOX}$ $EM_1$ and $T_{TOX}$ $EM_3$ cells with clone size samller (B) or greater / equal 3 (C). Dots are jittered in x and y direction. Numbers indicate proportions of $CD4^+$, $CD8^+$, $CD4^+$ $CD8^+$, or $CD4^-$ $CD8^-$ T-cells. D) TCR diversity was esimated for each LN patient sample using a rarefraction analysis (D). E-F) TCR diversity is exemplarily illustrated showing the 10 % most abundant clonotypes per sample. TCR: T-cell receptor.

### Supplementary Figure 6

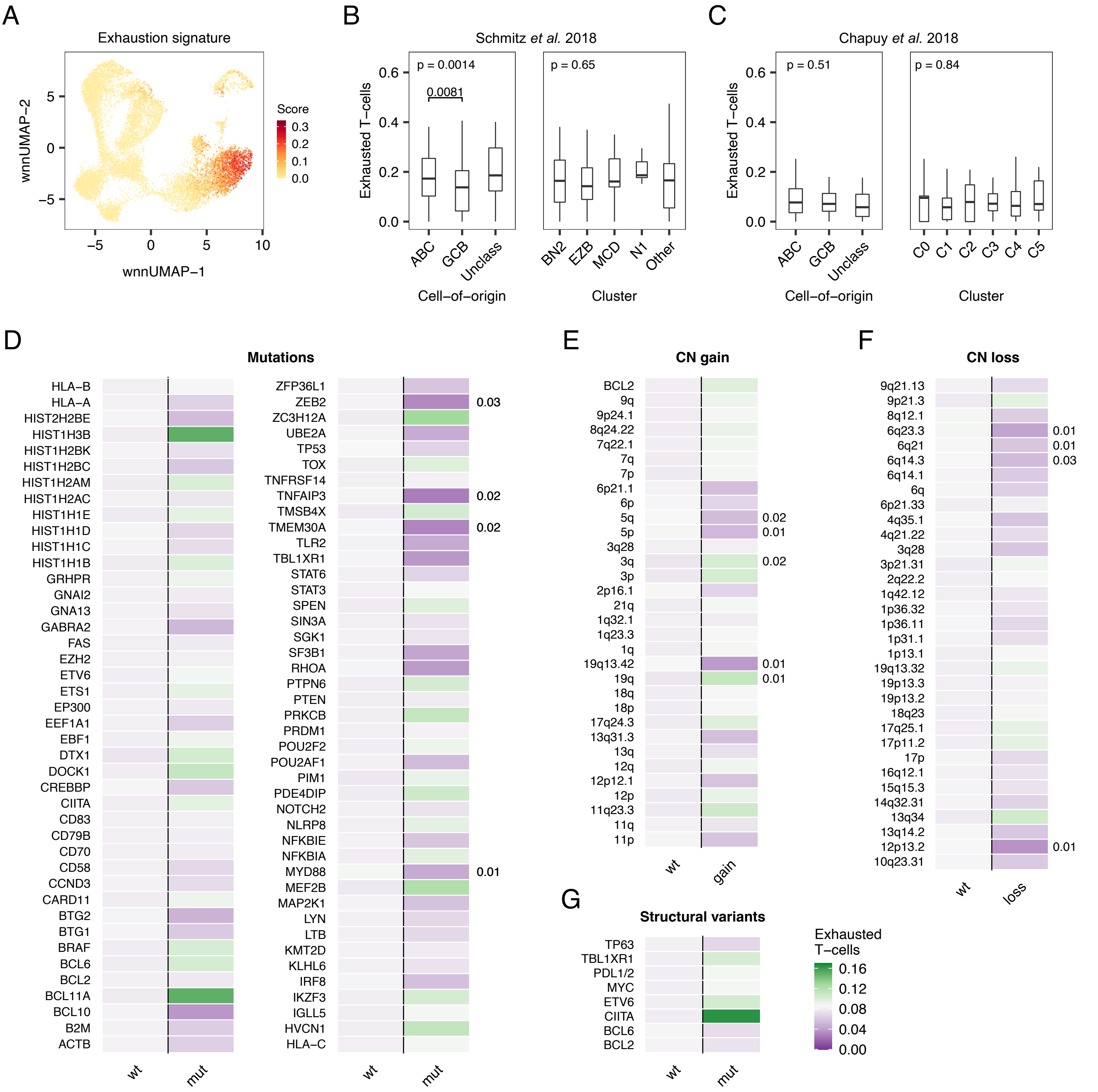


**Supplementary Figure 6.**

A) An exhaustion score was calculated based on a transcriptional signature of terminally exhaustion T-cells (see Method section for details) and then projected onto the reference UMAP plot. B-G) Bulk RNA-seq data from DLBCL patients from the Schmitz et al. cohort (B, n = 190) or the Chapuy et al. cohort (C-G, n = 137) were deconvoluted based on a gene expression signature of terminally exhausted T-cells. Box plots (B-C) show associations between T-cell exhaustion and cell-of-origin or genetic subtypes. Conditions were tested for significance using the Kruskal-Wallis-test. Wilcoxon-test was used as post-hoc test (B). Heatmaps (D-G) illustrate the mean estimated proportion of exhausted T-cells for somatic variants (D), amplifications (E), deletions (F), or structural variants (G), as indicated, based on deconvoluation of the bulk RNA-seq of the Chapuy et al. cohort (n = 137). Conditions were tested for significance using the Wilcoxon-test and corrected for multiple testing using the Benjamini-Hochberg procedure. After correction, all p values were > 0.05. To highlight the strongest differences, uncorrected p values ≤ 0.05 are shown to the right of each panel. ABC: Activated B-cell subtype. GCB: Germinal center B-cell subtype. CN: Copy number.

### Supplementary Figure 7

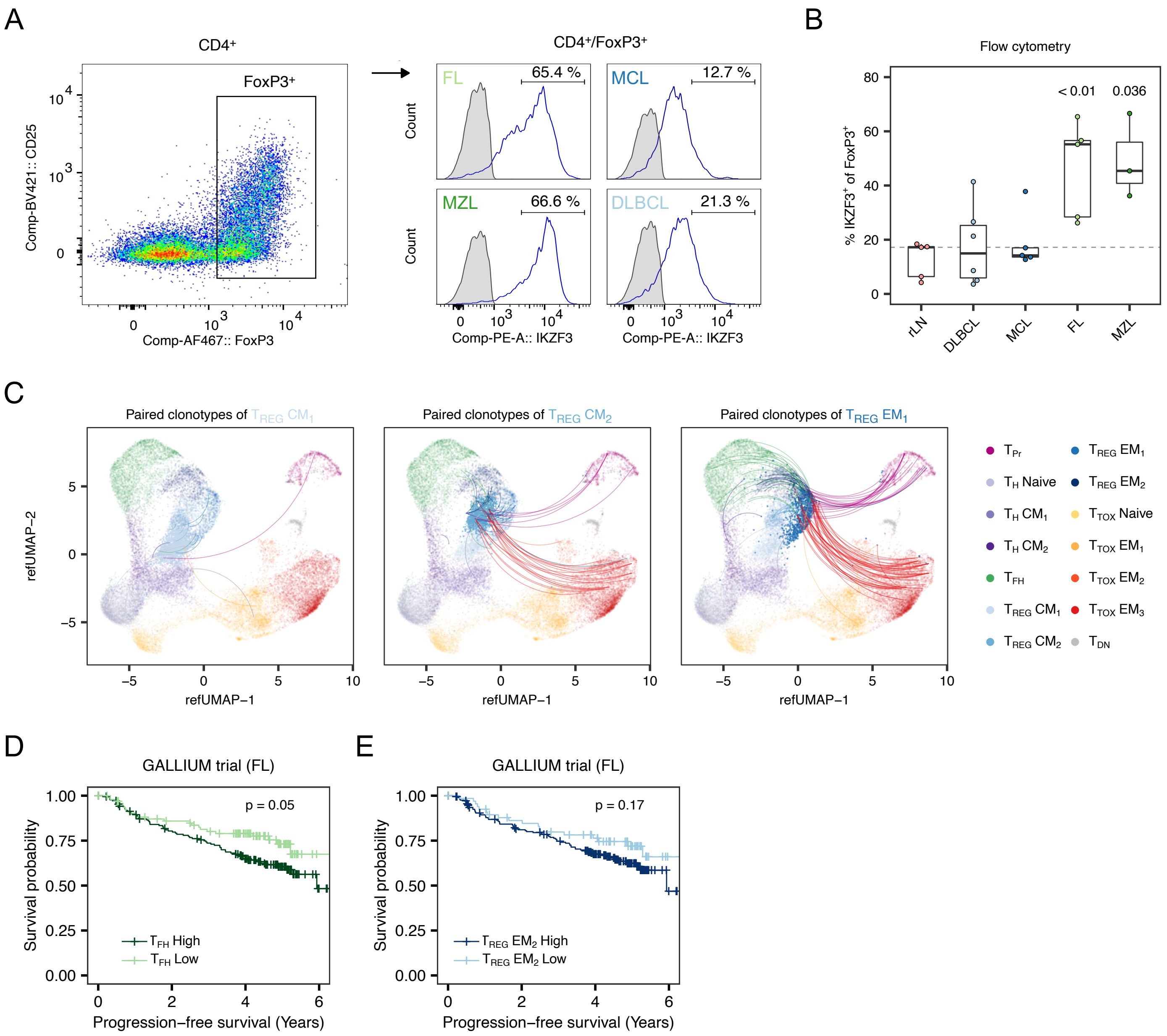


**Supplementary Figure 7.**

A) Representative pseudocolor and density plots showing the IKZF3 (Aiolos) protein expression in CD4+ FoxP3+ cells determined by flow cytometry. Numbers indicate percentage of positive cells based on gates as inidicated. Grey shaded histogramm represent fluorescence-minus-one control. B) IKZF3 protein expression in FoxP3+ T-cells determined by flow cytometry in 24 biologically independent B-cell lymphoma patient samples. Each entity was tested versus tumour-free samples (rLN) using the Wilcoxon-test. Shown are only p values $< 0.05$. C) 5’ scRNA alongside full-length TCR repertoire data from 11 biologically independent samples were mapped to the CITE-seq reference data. Lines connect all $T_{REG}$ $CM_1$, $T_{REG}$ $CM_2$ or $T_{REG}$ $EM_1$ cells with any other cell given that both T-cells have the same TCR clonotype. D-E) Bulk RNA-seq data from patients with FL were deconvoluted based on gene expression signature of $T_{REG}$ $EM_2$ (D) or $T_{FH}$ cells (E). Kaplan-Meier plots with p values of corresponding log-rank test. TCR: T-cell receptor.

### Supplementary Figure 8

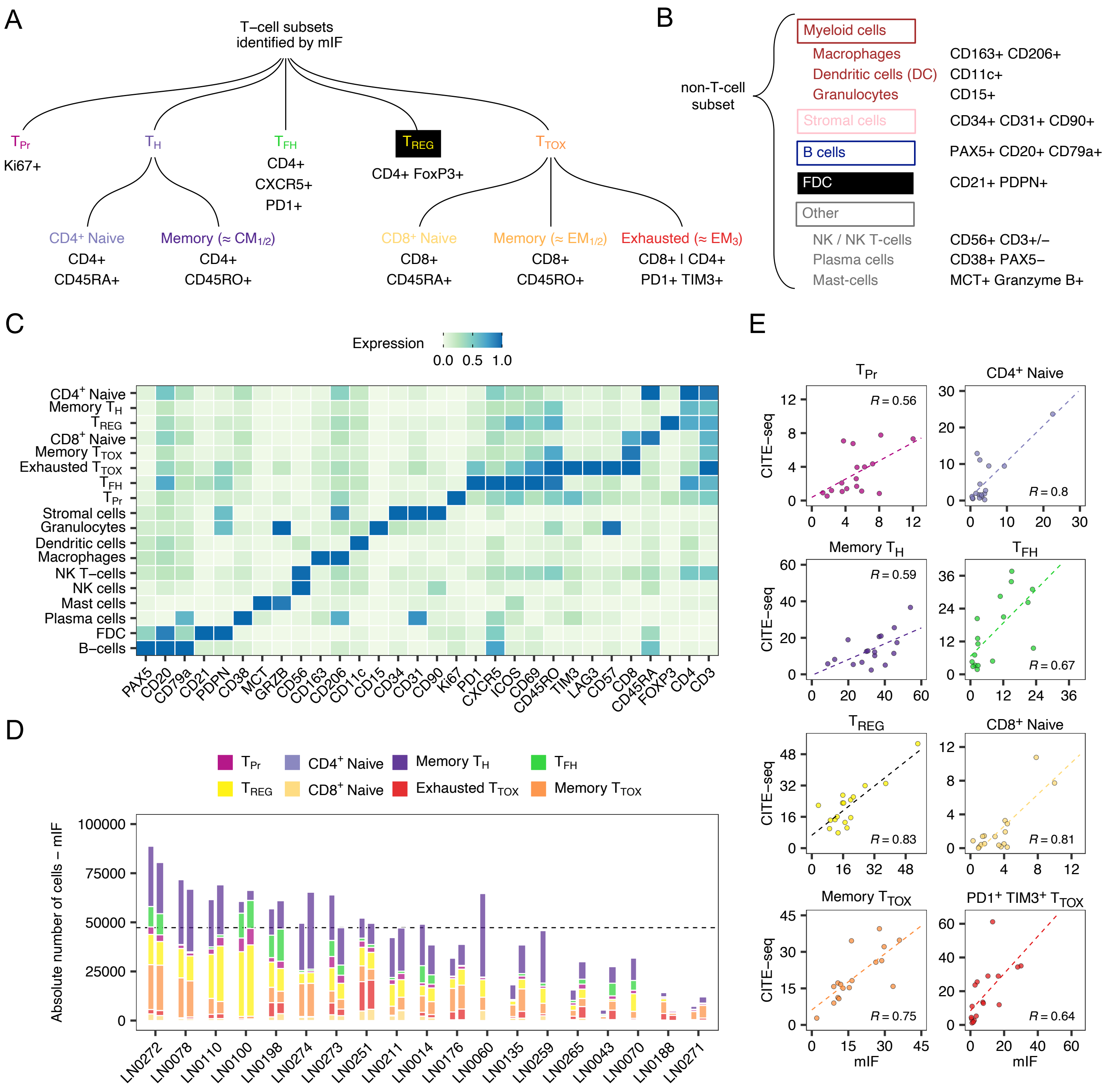


**Supplementary Figure 8.**

A-B) Overview of T-cell subsets (A) and other cell types (B) identified by highly multiplexed immunofluorescence including their marker profiles. C) Heatmap illustrating the mean expression of key marker proteins that were used to annotate the 18 cell types and T-cell subsets using highly multiplexed immunofluorescence. Expression values were scaled between 0 and 1. D) Shown are the absolute numbers of T-cells per subset, tissue core, and patient. A maximum number of two cores per patient were imaged. E) The low-granularity T-cell subpopulations detected by multiplexed immunofluorescence were aligned with the 14 high-granularity T-cell subsets identified by CITE-seq. Shown is the correlation of proportions for each of the T-cell subpopulations across the 19 LN samples that were analyzed by both approaches. Pearson's correlation coefficient is given for each panel (R). mIF: Multiplexed immunofluorescence.

### Supplementary Figure 9

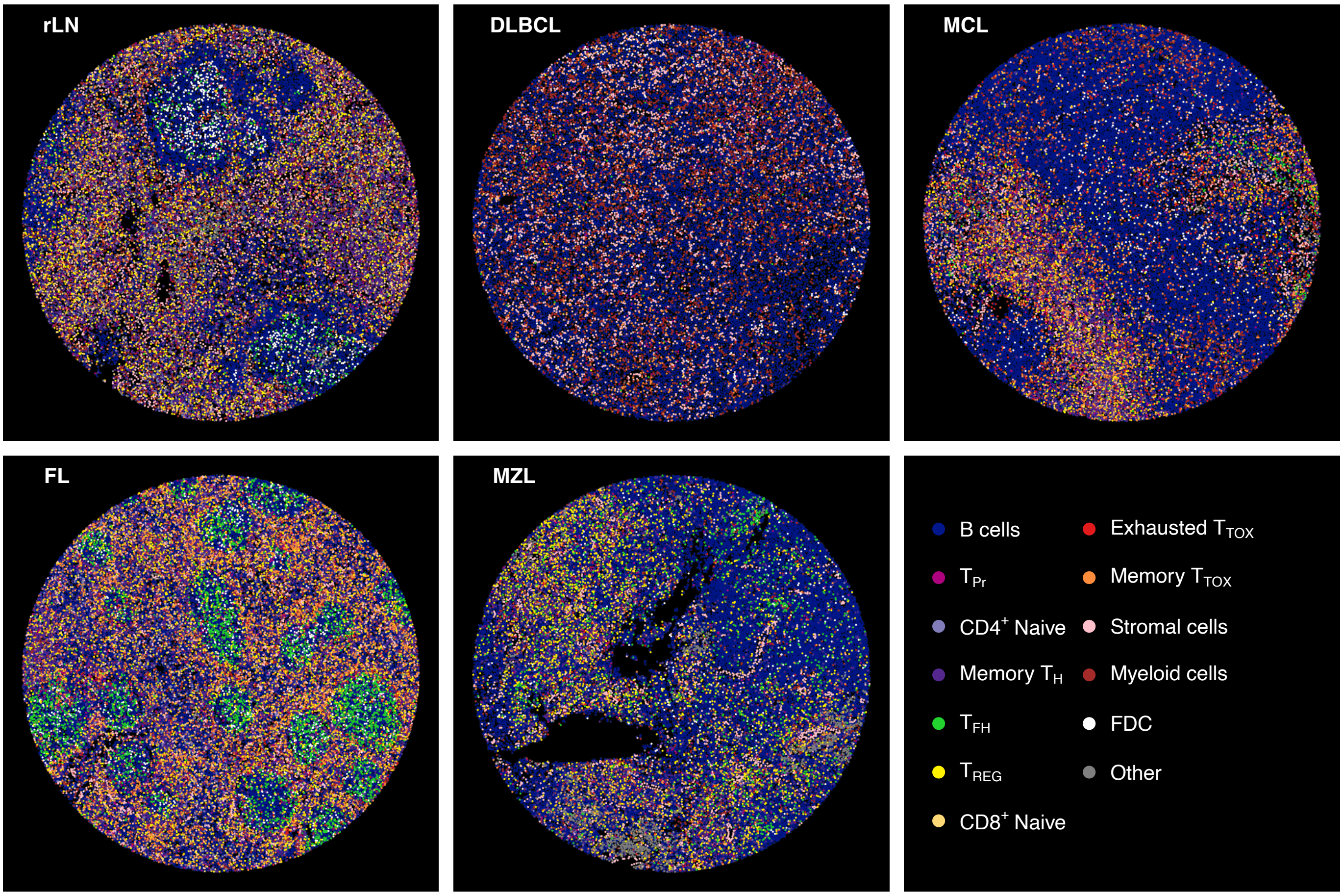


**Supplementary Figure 9.**
Shown are representative LN tissue cores for all entities investigated. Each cell was colored by T-cell subsets or non T-cell cell types as indicated. FDC: Follicular dendritic cell.

### Supplementary Figure 10

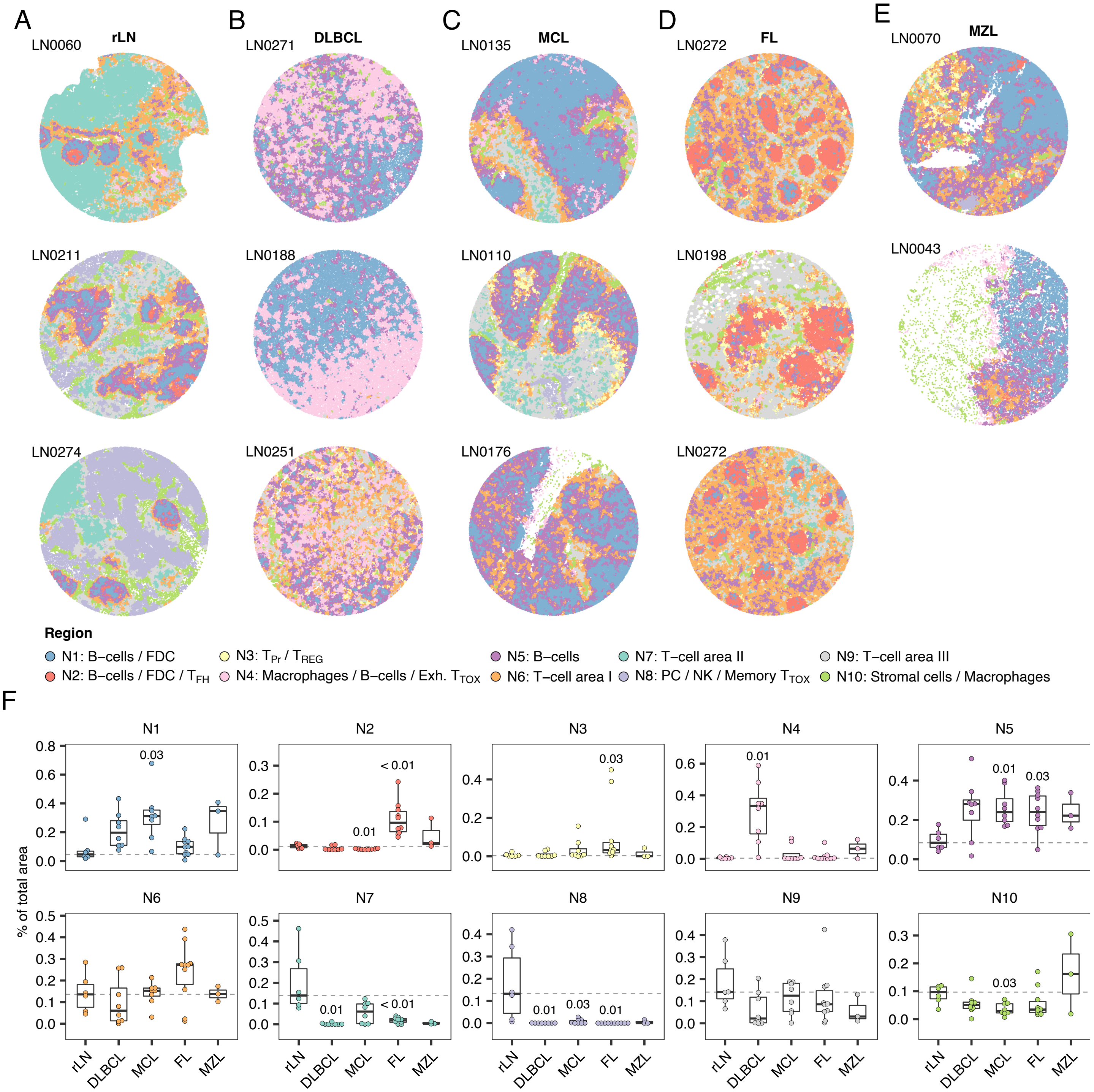


**Supplementary Figure 10.**

A-E) Representative LN-derived tissue cores from three or two different patients per entity. Each cell was colored according to its neighbourhood. F) Box plots showing the proportions of selected neighborhoods of each tissue core across n = 35 tissue cores. Each entity and neighborhood was tested versus rLN using the Wilcoxon-test. P values were corrected for multiple testing using the Benjamini-Hochberg procedure. Dashed lines indidcate the median of rLN. LN: Lymph node.
